# The Glyco-enzyme adaptor GOLPH3 Links Intra-Golgi Transport Dynamics to Glycosylation Patterns and Cell Proliferation

**DOI:** 10.1101/870477

**Authors:** Riccardo Rizzo, Domenico Russo, Kazuo Kurokawa, Pranoy Sahu, Bernadette Lombardi, Domenico Supino, Mikhail Zhukovsky, Anthony Vocat, Prathyush Pothukuchi, Vidya Kunnathully, Laura Capolupo, Gaelle Boncompain, Carlo Vitagliano, Federica Zito Marino, Gabriella Aquino, Daniela Montariello, Petra Henklein, Luigi Mandrich, Gerardo Botti, Henrik Clausen, Ulla Mandel, Toshiyuki Yamaji, Kentaro Hanada, Alfredo Budillon, Franck Perez, Seetharaman Parashuraman, Yusuf A Hannun, Akihiko Nakano, Daniela Corda, Giovanni D’Angelo, Alberto Luini

## Abstract

Glycans are ubiquitous sugar polymers with major biological functions that are assembled by glyco-enzymes onto cargo molecules during their transport through the Golgi complex. How the Golgi determines glycan assembly is poorly understood. By relying on the Golgi cisternal maturation model and using the glyco-enzyme adaptor and oncoprotein GOLPH3 as a molecular tool, we define the first example of how the Golgi controls glycosylation and associated cell functions. GOLPH3, acting as a component of the cisternal maturation mechanism, selectively binds and recycles a subset of glyco-enzymes of the glycosphingolipid synthetic pathway, hinders their escape to the lysosomes and hence increases their levels through a novel lysosomal degradation-regulated mechanism. This enhances the production of specific growth-inducing glycosphingolipids and reprograms the glycosphingolipid pathway to potentiate mitogenic signaling and cell proliferation. These findings unravel unforeseen organizing principles of Golgi-dependent glycosylation and delineate a paradigm for glycan assembly by the Golgi transport mechanisms. Moreover, they indicate a new role of cisternal maturation as a regulator of glycosylation, and outline a novel mechanism of action for GOLPH3-induced proliferation.

## Introduction

An important question in the fields of membrane transport and glycobiology is how glycans are assembled on protein and lipid scaffolds during transport through the Golgi complex. Indeed, most glycosylation reactions take place in the Golgi, which is often defined the ‘glycosylation factory’ of the cell. Glycans are ubiquitous sugar polymers that influence most cellular processes including adhesion, motility, signaling, and differentiation (Cummings, 2019; Stanley, 2016; Varki and Gagneux, 2015) and play major roles in several human diseases including cancer (Pinho and Reis, 2015; Stanley, 2011; Stowell et al., 2015). It would thus be of great value for biology and medicine to understand the mechanisms of glycan assembly in the Golgi and to clarify how these mechanisms affect bodily functions.

Glycosylation consists of orderly series of reactions catalyzed by glyco-enzymes (glycosyltransferases and glycosidases) (Colley et al., 2015; Stanley, 2011) organized in glycosylation pathways [*i.e*., the N- and O-linked glycosylation, the glycosphingolipid (GSL) pathway and the proteoglycan synthetic cascade (Stanley, 2011)]. The type and abundance of the glycans produced depend mainly on two factors: a) the gene expression pattern of the glyco-enzymes and accessory proteins (*e.g*., sugar and ion transporters) (Neelamegham and Mahal, 2016; Pothukuchi et al., 2019) and b) the order and frequency of the encounters between glyco-enzymes and substrates during transit through the Golgi stack (Bard and Chia, 2016; Pothukuchi et al., 2019; Stanley, 2011). The latter process, in turn, depends on the organization and dynamics of intra-Golgi transport and hence on the Golgi traffic machinery. Unfortunately, while significant progress has been made towards understanding the transcriptional programs that regulate glyco-enzymes expression and their effects on glycan structures (Neelamegham and Mahal, 2016; Varki and Gagneux, 2015), the mechanisms by which the intra-Golgi transport machinery controls glycosylation remain poorly understood.

This study capitalizes on recent advances in our comprehension of intra-Golgi traffic to unravel the links among the intra-Golgi transport machinery, glycosylation and the associated cell function. intra-Golgi traffic, although it depends on ‘canonical’ dissociative transport events [such as vesicle budding, docking, fusion (Glick and Nakano, 2009) *etc.*], is organized in a specific configuration called cisternal progression/maturation that differs from the classical linear anterograde vesicular transport mechanism (Emr et al., 2009; Glick and Nakano, 2009). The Golgi cisternal maturation is therefore likely to provide the correct framework to analyze the glycosylation process.

The Golgi comprises few stacked *cisternae* (*cis*, medial and *trans*) that contain glyco-enzymes distributed in a *cis-trans* order that roughly reflects the sequence of the reactions involved in the various glycosylation pathways (Stanley, 2011). Cargo proteins reach the *cis* Golgi from the endoplasmic reticulum (ER) within membranous carriers, which fuse homotypically to form new *cis cisternae*. These *cisternae* then convert into medial and *trans* cisternal elements through glyco-enzyme recycling, and finally disassemble into carriers destined to the plasma membrane (PM) or other destinations (cisternal progression) (Emr et al., 2009; Glick and Nakano, 2009). Glyco-enzymes recycle backward within COPI coated vesicles in synchrony with cisternal progression to maintain the compositional homeostasis of the progressing *cisternae* (cisternal maturation) (see scheme in **Fig. S1A** for a detailed description of the organization and mechanisms of cisternal maturation) (Bonifacino and Glick, 2004; Goud et al., 2018; Ishii et al., 2016; Jackson and Bouvet, 2014; Papanikou and Glick, 2014; Willett et al., 2013; Witkos et al., 2019; Wong and Munro, 2014). In this context, the cargo moving forward is sequentially intercepted by the retrogradely moving glyco-enzymes, resulting in glycan assembly. Glyco-enzyme recycling is therefore a key process in the regulation of glycosylation during intra-Golgi transport (**Fig. S1A**).

The recycling of glyco-enzymes requires specific adaptors that gate their entry into COPI coated retrograde transport vesicles. Among the adaptors so far identified the COPI δ and ς subunits and the Golgi phosphoprotein 3 (GOLPH3) are the best characterized (Casler et al., 2019; Liu et al., 2018; Schmitz et al., 2008; Tu et al., 2008). The role of GOLPH3 in enzyme recycling was first demonstrated in yeast by Banfield and Burd (Schmitz et al., 2008; Tu et al., 2008) and later shown to be conserved in *metazoa* including mammals (Ali et al., 2012; Chang et al., 2013; Eckert et al., 2014; Isaji et al., 2014; Pereira et al., 2014). The GOLPH3 yeast homologue (Vps74p) binds a specific motif in the cytosolic tail of a set of Golgi mannosyltransfersases (Vps74p clients) as well as the COPI coatomer. This results in the incorporation of the Vps74p clients into COPI recycling vesicles and hence in their retrograde transport within the secretory pathway. Thus, changing the levels of Vps74p alters the distribution of the Vps74p clients across the transport stations (Casler et al., 2019; Liu et al., 2018; Schmitz et al., 2008; Tu et al., 2008; Wood et al., 2009). Importantly, the human *GOLPH3* gene is amplified in solid tumors where it stimulates mitogenic signaling and cell proliferation, acting as an oncogene (Scott et al., 2009).

We have used GOLPH3 as a tool to manipulate Golgi enzyme recycling, and analyzed the entire sequence of events that lead from GOLPH3 binding to client glyco-enzymes to specific glycosylation patterns to mitogenic signaling and cell proliferation. We find that GOLPH3, acting as an integral component of the cisternal maturation mechanisms, selectively binds and retrotransports a specific subset of glyco-enzymes. GOLPH3-driven recycling, through an articulated series of mechanistic steps, promotes the increase of the levels of GOLPH3 clients. Among these, a functionally coherent group is placed at critical bifurcation points in the GSL metabolic network, so that their increase reprograms the GSL pathway towards overproduction of growth-promoting metabolites, resulting in enhanced mitogenic signaling and cell growth.

These findings define the first example of how the intra-Golgi dynamics control glycosylation and the associated cell functions. They also introduce a new view of cisternal maturation by which this mechanism operates as an assembly of recycling mechanisms acting on distinct enzymatic modules rather than as a simple constitutive housekeeping process. Moreover, they have broad pathophysiological and medical implications as they outline a novel mechanism of action for GOLPH3-induced proliferation based on GSL biosynthesis.

## Results

In order to investigate the role of GOLPH3 in Golgi glycosylation we started with the identification of mammalian enzymes that interact with GOLPH3.

### GOLPH3 binds and retro-transports a set of GSL synthetic enzymes by interacting with a novel motif in their cytosolic tails

To identify mammalian GOLPH3 clients, we used the knowledge available on the yeast GOLPH3 homologue Vps74p. Vps74p acts by binding to the consensus sequence (F/L)-(L/I/V)-x-x-(R/K) in the cytosolic tail of a set of glyco-enzymes (Tu et al., 2008). Since the two mammalian GOLPH3 isoforms (*i.e*., GOLPH3 and GOLPH3-like [GOLPH3L]) can partially replace Vps74p in phenotypic assays (though the former with lower efficiency than the latter) (Tu et al., 2008), it is likely that they also bind to the same consensus sequence in Vps74p clients. We thus asked whether GOLPH3 binds a similar motif also in mammalian enzymes (Tu et al., 2008). Examining the known direct GOLPH3 interactors in mammals [*i.e*., ST6GAL1 (Eckert et al., 2014), PomGNT1 (Pereira et al., 2014); and GCNT1 (Ali et al., 2012)] we found that they lack the complete yeast consensus motif, however they bear a similar sequence where the first hydrophobic amino acid is not present and the second is consistently a leucine [*i.e*., L-x-x-(R/K), for brevity here referred to as LxxR]. We thus looked for the LxxR sequence in type II human transmembrane glyco-enzymes and found it in the tails of 47 enzymes. Among these, enzymes of the GSL biosynthetic pathway are the most represented (with 19 glyco-enzymes accounting for 47.5% of the whole GSL synthetic pathway), followed by proteoglycan synthetic enzymes (13 glyco-enzymes; 22.8 % of the pathway) and by smaller groups of O- and N-linked glycosylation enzymes (11 and 8 glyco-enzymes; 23.4 and 22.2 % of the respective pathways) (**Fig. S1B**). Although arginines are very common in the glyco-enzyme tails, the LxxR motif is specifically enriched in GSL synthetic enzymes, indicating that this sequence likely is functionally significant. These enzymes are potential interactors of either GOLPH3, GOLPH3L, or both. In the following we focus on GOLPH3. A study on GOLPH3L will be presented separately.

Seeking functional testing of these conclusions with an independent approach, we overexpressed GOLPH3 (GOLPH3 OE) in HeLa cells and used a panel of lectins and bacterial toxins to detect changes in major glycosylation pathways by cytofluorimetry. As shown in **Fig. S1C**, GOLPH3 OE increased the levels of the GSL Gb3 (as assessed by Shiga toxin [ShTxB] staining) (Jacewicz et al., 1986) and, to a minor extent, of sialylated glycans at the cell surface (see **Fig. S1C** and methods for experimental details), while it had no detectable effects on other major glycosylation pathways. These data suggest that GOLPH3 fosters sphingolipid glycosylation possibly by interacting with GSL metabolic enzymes. We thus tested whether GSL enzymes that bear the LxxR motif bind GOLPH3. We used biotinylated peptides corresponding to the cytosolic tails of GSL enzymes in co-precipitation and isothermal titration calorimetry (ITC) experiments. We found that while the enzymes devoid of the LxxR motif fail to interact with GOLPH3, most of those that have LxxR show interaction with GOLPH3 (**Fig. 1A, B**) the exception being GA2/GM2 synthase (GM2S). This suggests that the LxxR motif contributes to, but it is not sufficient for, GOLPH3 binding.

**Figure 1.**
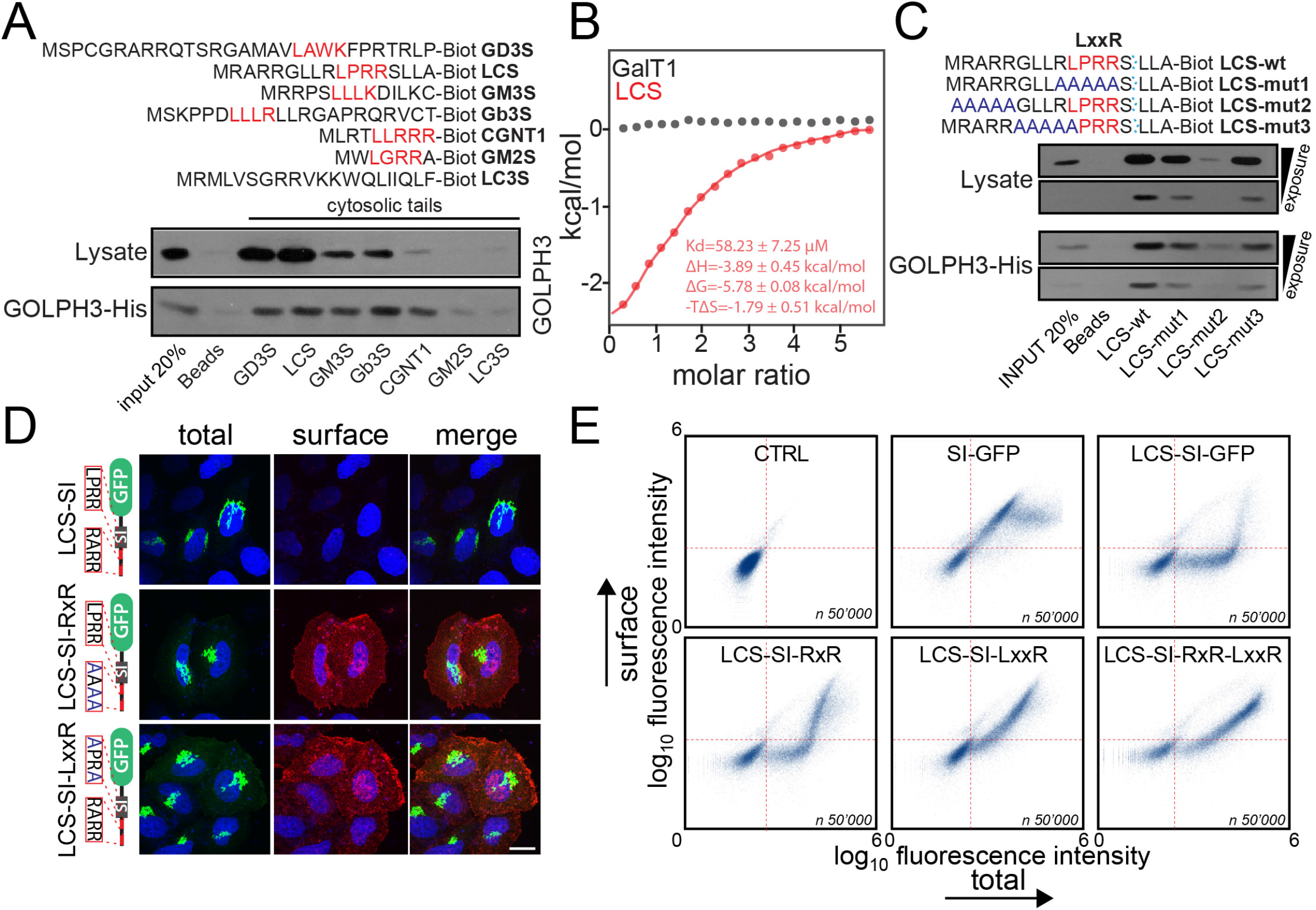
GOLPH3 interacts with the cytosolic tails of selected GSL synthetic enzymes. (A) Pull-down experiments involving biotinylated peptides of glyco-enzyme cytosolic tails from HeLa cells lysates (top blot) or purified His-tagged GOLPH3 protein (bottom blot). The N-terminal cytosolic tail of the involved glyco-enzymes is shown on top of the blot (LxxR motif in red, biotin at the C-terminus). Immunoblotting shows affinity-captured GOLPH3 (endogenous and recombinant). (B) ITC profile for biotinylated LCS and GalT1 cytosolic tails with recombinant GOLPH3. (C) Pull-down experiments of biotinylated LCS WT and LCS mutant cytosolic tails from HeLa cells lysates (top blot) or purified His-tagged GOLPH3 protein (bottom blot). The N-terminal cytosolic tail of LCS WT and LCS mutants (mut. 1, 2 and 3) is shown on top of the blot (mutations are in blue, LxxR motif in red, biotin at the C-terminus). (D) HeLa cells expressing luminal SI-GFP chimeras of sucrose-isomaltase (SI-GFP, green) containing the WT and mutant N-terminal cytosolic tails of LCS, were fixed and labelled for surface localized GFP (red). Cells expressing LCS-SI-RxR and LCS-SI LxxR proteins shown PM staining. Bar, 20 μm. (E) Quantitative flow cytometry-based analysis of the experiment reported in D; n is indicated in the graph.

To identify the complete GOLPH3 recognition motif, we focused on lactosylceramide (LacCer) synthase (LCS; the best binder in the above co-precipitation assays that we also confirm to co-immunoprecipitate GOLPH3 when expressed as a full-length protein in HeLa cells) (**Fig. 1A-B** and **Fig. S1D**). We introduced mutations in LCS and tested the mutants for GOLPH3 binding (**Fig. 1C**). Mutating either the LxxR containing region, or a positively charged cluster upstream of LxxR, caused binding inhibition, indicating that GOLPH3 interacts with LCS tail through both of these regions. We thus re-examined the cytosolic tails found by us or others to interact with GOLPH3 (above) and noticed that they all exhibit at least one positive charge 3 to 6 amino acids upstream of the LxxR motif. We call this motif (H/R/K [x]_2-5_ LxxR/K) the dual GOLPH3 binding (DGB) motif. We then searched for the DGB motif in glyco-enzyme tails and found it in 23 enzymes, 9 of which belong to the GSL metabolic pathway while others are distributed among proteoglycan (4 enzymes), N-linked (3 enzymes) and O-linked (5 enzymes) glycosylation pathways (**Fig. S1E, F**). Notably, the frequency of the DGB motif is twice higher in the GSL than in the other glycosylation pathways. To confirm this motif, we tested 3 additional DGB motif bearing tails (FUT5, B3GNT4, and GALNT3) for interaction with purified recombinant GOLPH3. As shown in **Fig. S1G**, peptides corresponding to the cytosolic tails of these enzymes interacted with recombinant GOLPH3 *in vitro*. Altogether, the 10 DGB-bearing tails tested by us or others did bind GOLPH3, while none of the 8 tails devoid of the motif was able to bind this adaptor (**Fig. S1H**) indicating that the presence of DGB motif reliably predicts binding to GOLPH3. These data, however, do not formally exclude that further (so far undetected) variants of the LxxR motif or even unrelated motifs might result in efficient GOLPH3 binding, and a dedicated structural study involving the identification of the GOLPH3 binding site/s for its clients is required to unequivocally solve this issue. Moreover, they do not necessarily imply that the DGB motif supports GOLPH3-based Golgi retention of all DGB-bearing enzymes in live cells. We call these enzymes putative GOLPH3 clients.

We next examined whether the DGB motif is sufficient for retention in the Golgi in live cells. We expressed a chimeric protein comprising the LCS cytosolic tail at the C-terminus of the transmembrane domain (TMD) of sucrose isomaltase (SI), a type II integral PM protein, fused with GFP at the N-terminus [LCS-SI-GFP,(Liu et al., 2018)] in HeLa cells. This reporter does not have Golgi retention signals in its TMD and luminal domains, thus its retention would be ascribed exclusively to the enzyme tail (Liu et al., 2018). As shown in **Fig. 1D, E**, the LCS tail retained the SI construct at the Golgi throughout a wide range of expression levels and only cells that overtly overexpress it showed the reporter at the PM, indicating that the LCS tail mediates Golgi retention by a saturable mechanism. Mutating the LxxR sequence or the upstream positively charged residues or both, resulted in loss of Golgi localization (**Fig 1D, E**) indicating that the DGB motif in the LCS tail is responsible for retention of these constructs, most likely by mediating their interaction with GOLPH3; and suggesting that GOLPH3 is required for the Golgi retention of these glyco-enzymes. Indeed, GOLPH3 depletion markedly inhibits the Golgi retention of constructs bearing the native tail peptides of LCS, GM3 synthase (GM3S), Gb3 synthase (Gb3S), and GD3 synthase (GD3S). Further experiments with full-length (TMD-containing) LCS and other GSL enzymes based on assays in cells depleted of GOLPH3 confirmed that GOLPH3 is required for Golgi retention of glyco-enzymes that bear the DGB motif and does not affect retention of all the DGB-devoid enzymes tested in agreement with the *in vitro* binding data (see below). We define these GOLPH3-retained enzymes verified GOLPH3 clients.

Notably, one of the tested GOLPH3 clients, Gb3S, has been shown to require the interaction with the nonaspanin TM9SF2, possibly through its TMD, for retention in the Golgi (Yamaji et al., 2019); other enzymes including LCS exhibited a similar behavior. We reproduced these data under our conditions (not shown) which suggests that at least some of the DGB motif bearing enzymes are retained in the Golgi through two different domains, one that binds GOLPH3 in the cytosolic tail, and the other in the TMD that interacts with TM9SF2. A complete dissection of the retention mechanism of these enzymes is beyond the scope of this paper and will be reported separately.

In sum, GOLPH3 plays a necessary though not exclusive role in the Golgi retention of several glyco-enzymes through binding to the DGB motif in their cytosolic tails. Here, we focus on the GSL metabolic GOLPH3 client enzymes as several GSL metabolites possess growth-promoting bioactivity and might play a role in the oncogenic activity of GOLPH3 (Canals and Hannun, 2013; Furukawa et al., 2019; Morad and Cabot, 2013).

### GOLPH3 retains its client enzymes in the Golgi acting as a component of the cisternal maturation machinery

Based on the available knowledge on GOLPH3 mechanism of action, GOLPH3 retains its clients in the *trans* Golgi by promoting their inclusion into retrograde COPI vesicles and by recycling them backward towards medial *cisternae* (Eckert et al., 2014; Tu et al., 2008). This is the same mechanism proposed for enzyme recycling in cisternal maturation, which suggests that GOLPH3 operates in the context of the maturation mechanism. However, the observation that GOLPH3 acts on a selected set of glyco-enzymes is partially inconsistent with cisternal maturation that requires the synchronous recycling of all Golgi enzymes and residents to maintain the compositional homeostasis of Golgi *cisternae* (Glick and Luini, 2011; Nakano and Luini, 2010). Indeed, it has been proposed that GOLPH3 might mediate a salvage mechanism to retrieve selected enzymes that occasionally escape the Golgi stack, rather than participating in cisternal maturation. To distinguish between these two possibilities, we considered that cisternal maturational and salvage mechanisms must entail different kinetic features in their transport behaviors (see scheme in **Fig. S1A**).

In fact, if GOLPH3 operates in the context of maturational recycling, its removal should result in complete loss of clients retention in the Golgi, which should traverse the Golgi stack at exactly the same rate as secretory cargoes. Instead, if GOLPH3 is involved only in the salvage of escaped enzymes, its removal should result simply in the gradual and slow leakage of the clients out of the stack. We thus, designed experiments in both mammalian and yeast cells to test whether or not GOLPH3 (or Vps74p) recycles its client enzymes in synchrony with cisternal progression.

For mammalian cells, we examined the trafficking of the GOLPH3 client LCS, both by itself and compared to that of a cargo reporter; we used the HeLa-M strain that does not express GOLPH3L (our unpublished observation) in order to avoid possible interference from this adaptor that might interact with some of the GOLPH3 clients. We first synchronized the transport of LCS by using a LCS-GFP/ KDEL-hook encoding plasmid (LCS-GFP_RUSH; see **Methods** for a detailed description) (Boncompain et al., 2012), that can be arrested and accumulated in the ER; then we released this construct synchronously by adding a suitable ligand (Boncompain et al., 2012) and monitored its progression across the Golgi. In control cells, LCS-GFP_RUSH moved from *cis* to *trans*-Golgi in 10 to 20 min and then remained in the *trans*-Golgi, without reaching the TGN, for up to 2 hours. Conversely, in GOLPH3-KD cells, LCS-GFP_RUSH traversed the Golgi without stopping in the *trans cisterna* and reached the TGN in less than 20 min. This rate is typical of cargo transport by cisternal maturation (Mironov et al., 2001) (**Fig. 2A**).

**Figure 2.**
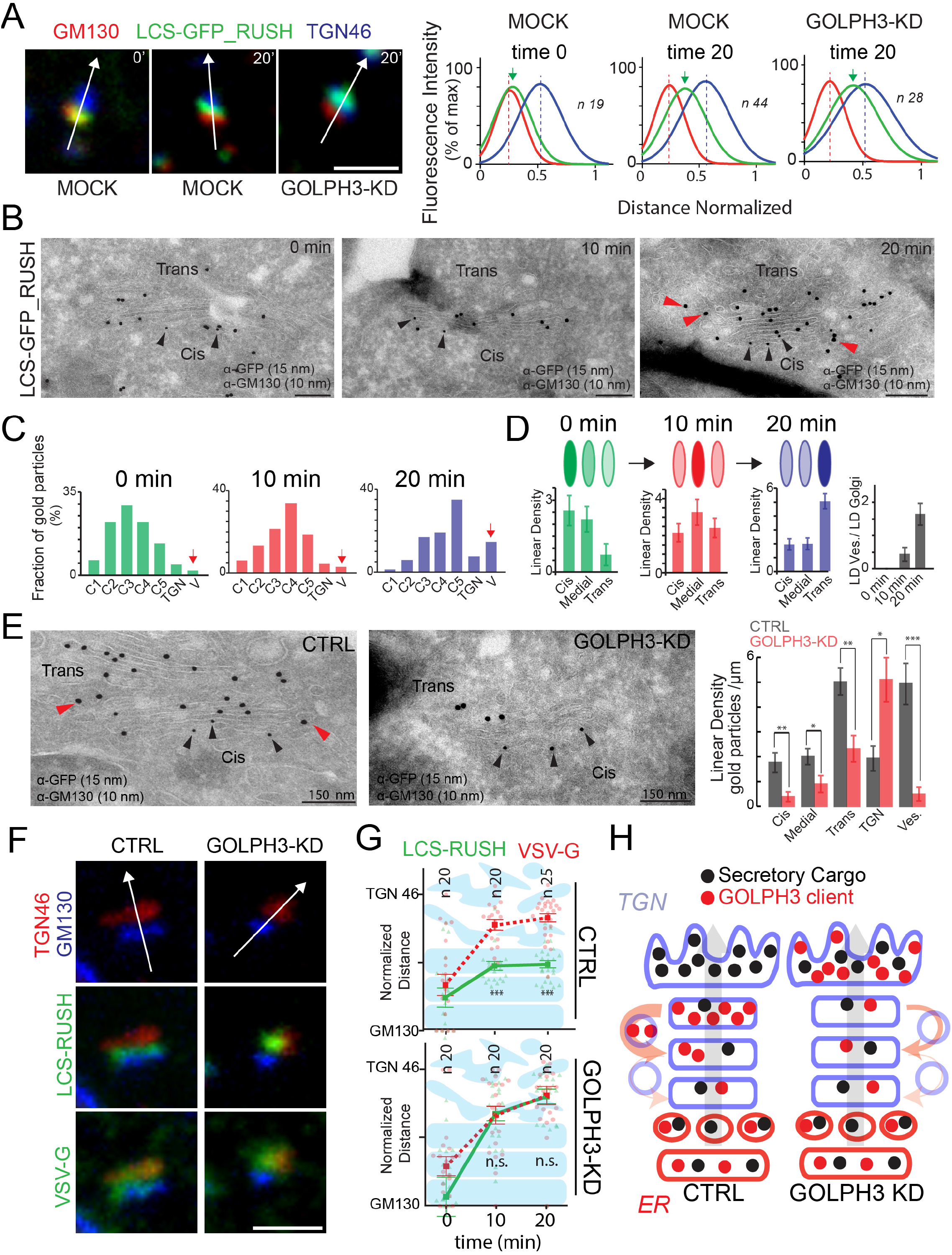
GOLPH3 drives the intra-Golgi recycling of LCS. (A) HeLa cells mock-treated or GOLPH3-KD were transiently transfected with LCS-GFP_RUSH and subjected to the synchronization protocol described in **Methods**. Cells were then subjected to IF: LCS-GFP_RUSH (green), *cis* Golgi marker GM130 (red), and TGN marker TGN46 (blue). Representative individual Golgi mini-stacks are shown. Bar, 1 μm. Quantification (right graph) by computational coalescence of line scans was performed by normalized line scan analysis (normalization of the distances was performed considering the GM130 peak as 0, and the TGN46 peak as 1). The n indicated in the graph refers to the number of Golgi stacks analysed in each condition. (B) HeLa cells were transfected with LCS-GFP_RUSH, subjected to the synchronization protocol (see **Methods**), fixed and processed for cryoimmunolabeling with an anti-GFP antibody (15 nm gold particles) and anti-GM130 antibody (10 nm gold particles, black arrowheads) peri-Golgi vesicles containing LCS-GFP_RUSH are marked by red arrowheads. Bar 150 nm. (C, D) Quantification of the distribution of LCS-GFP_RUSH as measured by Frequency Distribution Analysis (C) and linear density (LD, in D) (data are means ± SEM). (E) HeLa cells expressing LCS-GFP_RUSH mock or GOLPH3-KD were treated and stained as in B. Black arrowheads indicate GM130; peri-Golgi vesicles containing LCS-GFP_RUSH are marked by red arrowheads. Quantification of LCS-GFP-RUSH distribution (right graph, linear density, LD) across the stack and in peri-Golgi vesicles (data are means ± SEM; *p <0.05, **p < 0.01; ***p <0.001 [Student’s t-test]). Bar 150 nm. (F) Mock and GOLPH3-KD HeLa cells co-expressing LCS-GFP_RUSH and VSV-G were subjected to the co-synchronization protocol described in **Methods**. Cells were then subjected to IF: LCS-GFP-RUSH or VSV-G (green), GM130 (blue), and TGN46 (red). Representative individual Golgi mini-stacks are shown. Bar, 1 μm. (G) Quantification of the experiment in (F) was performed by normalized line scan analysis (normalization of the distances was performed considering the GM130 peak as 0, and the TGN46 peak as 1); n indicated in the graph refers to the number of Golgi stacks analysed in each condition, individual data points and means ± SEM are shown, ***p < 0.001 (Student’s t test); ns, not significant). (H) Model of the intra-Golgi transport of the GOLPH3 client enzyme (red dots) and of the secretory cargo protein (black dots) in the presence (left) or in the absence (right) of GOLPH3, through the stack. In control cells (left) the GOLPH3 client moves through the *cis* and medial *cisternae* according to the progression-maturation mechanism (see **Fig. S1A**) without entering peri-Golgi vesicles (hence, without leaving the lumen of the *cisternae*). When the GOLPH3 client reaches the *trans cisterna*, where it resides, it enters the peri-Golgi COPI vesicles and recycles backward (orange arrow) to the medial *cisternae*, which are maturing into *trans* elements, and is thus permanently retained within maturing *trans* Golgi *cisternae*. Secretory cargo in contrast, moves together with the GOLPH3 client through the *cis* medial *cisternae* but then continues to move forward into the TGN. In the absence of GOLPH3, the GOLPH3 client moves through the stack and reaches the TGN together with secretory cargo.

Cryo-immunoelectron microscopy (cryo-IEM) confirmed that in control cells, LCS-GFP_RUSH progresses from *cis* to medial and then *trans cisterna* in about 10 min and then it remains in the *trans cisterna* (**Fig. 2B-D**). Moreover, it showed that, during progression form the *cis* to the *trans cisternae* (first 10 min) LCS-GFP_RUSH is excluded from peri-Golgi vesicles, while, upon reaching the *trans cisterna* where it is retained, it enters these vesicles (**Fig. 2B-D**). In contrast, in GOLPH3-KD cells, LCS-GFP_RUSH was not retained in the *trans cisterna* and reached the TGN in less than 20 min, in agreement with the IF data; and was barely detected in peri-Golgi vesicles (**Fig. 2E**). These data are coherent with the notion that LCS traverses the Golgi stack by cisternal maturation without leaving the cisternal lumen, and that it is retained in the *trans cisterna* by GOLPH3-dependent recycling via incorporation into retrograde COPI coated vesicles.

Finally, we monitored the progression of LCS-GFP_RUSH and of the cargo reporter vesicular stomatitis virus G glycoprotein [VSV-G; that crosses the Golgi by cisternal maturation (Mironov et al., 2001)] through the stack in the same cells by co-expressing and cosynchronizing these two proteins. To synchronize VSV-G we exploited a ts045-VSV-G mutant that can be accumulated in the ER at 40 C and can then be released from the ER by shifting the temperature to 32 C (Mironov et al., 2001), and combined this with the RUSH procedure (for LCS-GFP_RUSH) to co-synchronize the two reporters (see **Methods** for details). In control cells, LCS-GFP_RUSH and VSV-G entered the Golgi and moved through the *cis*/medial Golgi compartments together; then, LCS-GFP_RUSH was retained in the *trans*-Golgi, while VSV-G crossed the stack and reached the TGN in 20 min (**Fig. 2F-H**). On the contrary, in GOLPH3-KD cells, LCS-GFP_RUSH was not retained in the *trans*-Golgi and reached the TGN at the same rate as VSV-G (**Fig. 2F-H**); thus, LCS-GFP_RUSH and VSV-G crossed the Golgi stack together. As a specificity control of the effects of GOLPH3-KD we examined the localization of GalT1, a trans-Golgi enzyme that does not bind GOLPH3 (see **Fig. 1B**) both in the presence and in the absence of GOLPH3. GalT1 remained in the *trans cisterna* under both conditions, supporting the selectivity of action of GOLPH3 on LCS trafficking (**Fig. S2A**).

For experiments in yeast, we monitored Mnn9p, a client of the GOLPH3 yeast homologue Vps74p (Tu et al., 2008). At steady state, Mnn9p populates the *cis-*Golgi in WT cells, while in *vps74Δ*, Mnn9p localizes in the vacuole and in the *trans-*Golgi (as marked by Sys1p) (**Fig. S2B**). After 60 min of protein synthesis inhibition, Mnn9p was specifically lost in *vps74Δ* cells (**Fig. S2C**) indicating that, as for LCS and GOLPH3, the retention and polarized localization of Mnn9p at the Golgi depends on Vps74p. When Mnn9p transport was visualized *in vivo* (Ishii et al., 2016; Matsuura-Tokita et al., 2006) we observed that, while in WT cells Mnn9p was recycled out of maturing *cis cisterna* as soon as it started acquiring the *trans* marker Sys1p (Losev et al., 2006; Matsuura-Tokita et al., 2006), in *vps74Δ* cells Mnn9p remained in the *cisterna* until it was filled with the Sys1p *trans* marker (**Fig. S2D**).

We then visualized Mnn9p moving synchronously through the secretory pathway using the superfolder-GFP tag (Mnn9p-sfGFP) (Pedelacq et al., 2006). We photobleached Mnn9p-sfGFP and monitored the newly synthesized Mnn9p-sfGFP as it progressed through the secretory pathway. In WT cells, newly synthesized Mnn9p-sfGFP was quickly transported (2-4 min) to the Golgi and then remained there for the duration of the experiment (*i.e*., up to 30 min), while in *vps74Δ* cells Mnn9p-sfGFP was not retained and proceeded to the vacuole within 5 min (**Fig. S2E**). Notably the observed traffic effects are not due to differences in the Mnn9p-sfGFP synthetic rates between the two yeast strains (**Fig. S2F**), and the transport rate of Mnn9p-sfGFP to the vacuole in *Vps74Δ* cells is similar to that of a vacuolar cargo protein (Vph1p-sfGFP) (**Fig. S2G**).

Altogether, the results in both mammalian and yeast cells indicate that GOLPH3 and Vps74p are responsible for the retention of their clients at the Golgi complex by recycling these enzymes in lockstep with the progression of cargo containing *cisternae*. Thus, GOLPH3 and Vps74p operate in manner that is kinetically fully consistent with the maturation process (but not with a salvage mechanism), indicating that they are components of the maturation mechanism. Yet, they act on a selected pool of enzymes. This is incompatible with the classical view of cisternal maturation as a homogeneous housekeeping process operating equally on all enzymes, and calls for a revision of the maturation model (see **Discussion**).

### GOLPH3 regulates the levels of its client enzymes via its recycling action, by reducing their escape to the lysosomes and hence their lysosomal degradation

We next examined whether GOLPH3 can affect the behavior of its clients through its recycling action. To modulate GOLPH3-mediated recycling we manipulated its expression levels, both in HeLa cells and primary human fibroblasts (PHFs), using LCS as a model client. In PHFs, GOLPH3 OE did not apparently change the Golgi localization of endogenous LCS. Unexpectedly, however, it markedly increased its levels over those of control cells. GOLPH3 KD had the opposite effects, *i.e*., it markedly reduced LCS levels (**Fig. 3A**). In HeLa cells, GOLPH3 OE induced phenotypes that were qualitatively similar to those seen in PHFs (*i.e*., it increased the endogenous LCS levels, **Fig. 3B**) while the impact of GOLPH3 KD on endogenous LCS was difficult to assess reliably because of the very low expression levels of this enzyme (a feature common to most glyco-enzymes). We thus performed similar experiments using HA-tagged LCS and found that in HeLa cells, GOLPH3 OE and KD had effects similar to those observed in PHFs on endogenous LCS. In addition, GOLPH3 OE partially redistributed LCS to the ER (**Fig. 3C**), reminiscent of analogous observations in yeast (Tu et al., 2008). Western blot experiments confirmed the IF data, *i.e*., that GOLPH3 OE increased, and GOLPH3 KD decreased, the LCS-HA protein levels in HeLa cells (**Fig. 3D**).

**Figure 3.**
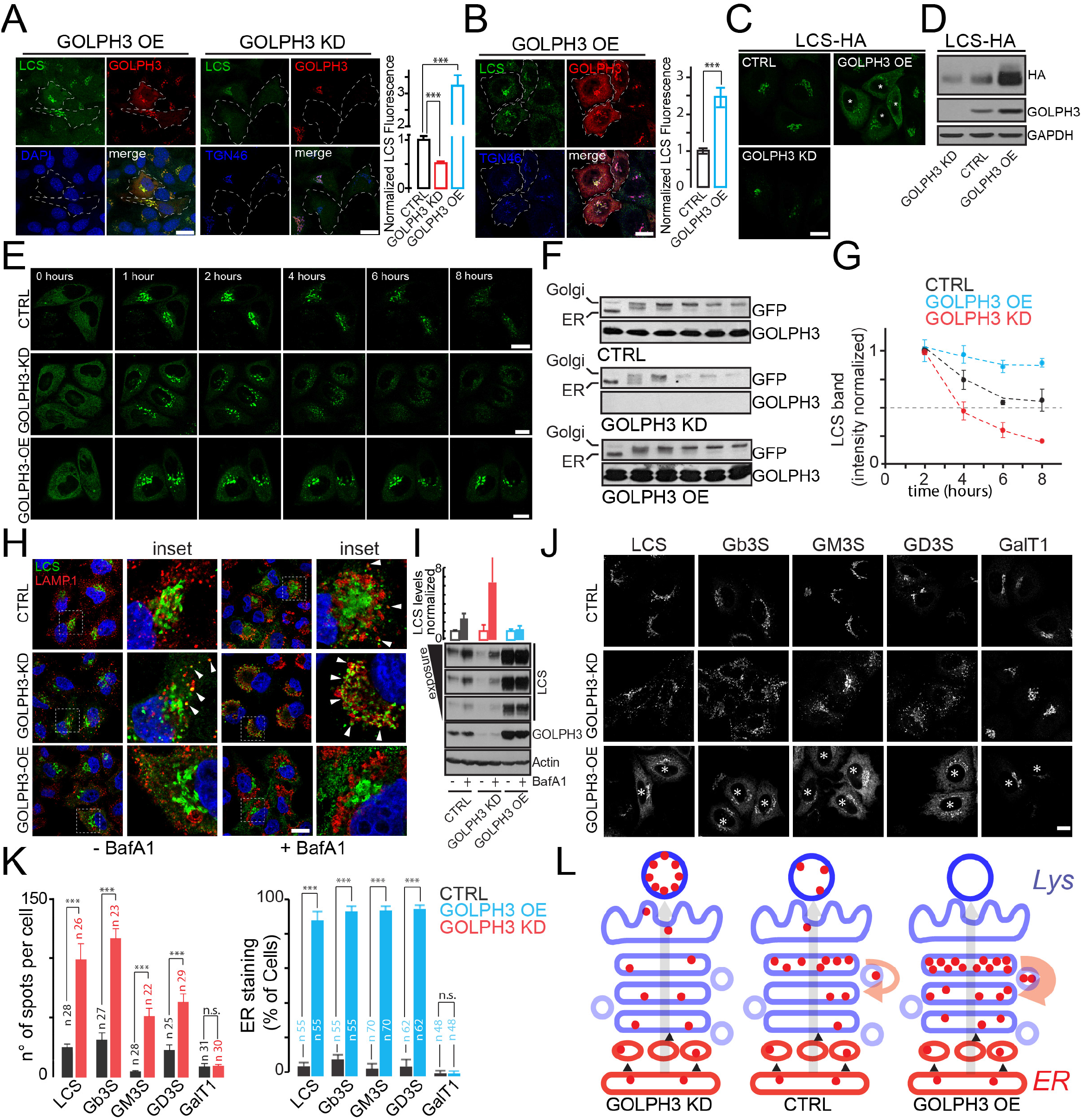
GOLPH3 regulates the lysosomal degradation of its clients. (A) PHFs GOLPH3 OE (left), or KD (right), were fixed and processed for IF: Endogenous LCS (green), GOLPH3 (red), TGN46 or DAPI (blue). Dashed lines indicate cells overexpressing or interfered for GOLPH3. Bar, 20 μm. Quantification of the amount of endogenous LCS (right graph) at the Golgi; data are means ± SEM; ***p <0.001). (B) GOLPH3-OE HeLa cells were fixed and processed for IF: Endogenous LCS (green), GOLPH3 (red), TGN46 (blue). Dashed lines indicate cells overexpressing GOLPH3. Bar, 20 μm. Quantification of the amount of endogenous LCS (right graph) at the Golgi (data are means ± SEM; ***p <0.001). (C) GOLPH3 OE or KD HeLa cells expressing LCS-HA were fixed and processed for IF: HA (green). Asterisks indicate GOLPH3 OE cells. Bar, 20 μm. (D) Representative WB of cells treated as in (C). (E) Assessment of LCS-GFP_RUSH synchronized transport in GOLPH3 OE or KD HeLa cells. Micrographs show cells at the indicated trafficking time points. Bar, 20 μm. (F) Representative WB of cells treated as in (E) (n = 3). (G) Plots of the densitometric quantification of the amount of LCS-GFP-RUSH (Golgi mature band) over time, starting from 2 hours chase (n = 3, data are means ± SD). (H) GOLPH3 OE or KD HeLa cells expressing LCS-HA were treated with bafilomycin A1 (BafA1, 10 nM) for 16 hr, fixed, and processed for IF labelling: LCS (green), LAMP1 (red), and DAPI (blue). Arrowheads indicate LCS and LAMP1 colocalization. Bar, 20 μm. (I) WB of cells treated as in (H). Densitometric quantification of the blot is plotted in the top panel (n = 3, data are means ± SD). (J) GOLPH3 OE or KD HeLa cells expressing the indicated HA-tagged glyco-enzymes were fixed and processed for IF. Asterisks indicate GOLPH3 OE cells. Bar, 20 μm. (K) Number of glyco-enzymes-HA positive spots per cell in GOLPH3-KD cells (left panel) and percentage of cells with glyco-enzymes-HA in the ER in GOLPH3-OE cells (right panel) are plotted (n is indicated in the graph; data are means ± SEM; ***p <0.001). (L) Regulation of the LCS levels by GOLPH3-dependent lysosomal degradation. In control cells (CTRL) LCS (red dots) progresses through the stack without leaving the cisternal lumen and without being captured in transport vesicles, according to the cisternal progression maturation mechanism (see **Fig. S1A**). When it reaches the *trans cisterna* it recycles backwards by GOLPH3 via COPI vesicles (thin orange arrow) and is therefore retained in this *cisterna.* A fraction of LCS reaches the TGN and then the lysosome, where it is degraded. Thus, LCS is degraded at a rate dependent on the efficiency of the GOLPH3-dependent recycling and retention action, or, in fact, on the competition between the GOLPH3 recycling mechanism and the rate of cisternal progression. In cells depleted of GOLPH3 (GOLPH3 KD), LCS is not retained in the stack and reaches the lysosome very rapidly. In cells overexpressing GOLPH3 (GOLPH3 OE) LCS reaches the *trans cisterna* as in control cells, but is then very efficiently retained in the Golgi stack by GOLPH3-dependent recycling (thick orange arrow). This prevents LCS from reaching the lysosomes, resulting in accumulation of LCS in the stack.

Cryo-IEM showed that, in control cells, LCS-HA displays a peak of concentration at the *trans*-Golgi, as expected, with lower levels in the TGN and in medial and *cis cisternae*, and is detectable in peri-Golgi vesicles, which were decorated by GOLPH3 (**Fig. S3A**). GOLPH3 OE caused a modest shift of LCS from *trans* towards early Golgi compartments, and increased LCS-HA in peri-Golgi vesicles. On the contrary, GOLPH3 KD resulted in low LCS-HA levels in the Golgi stack and in the absence of LCS-HA from vesicles (**Fig. S3A**). In sum, the OE of GOLPH3, that localizes at the *trans*-Golgi and at the TGN and in peri-Golgi vesicles (**Fig.S3B**), increases the levels of LCS and causes a moderate shift of this enzyme towards the medial and *cis*-Golgi, while LCS-HA leaves the stack rapidly in the absence of GOLPH3 (see below).

The effects of GOLPH3 OE on the levels of LCS could be either due to changes in LCS synthesis or in LCS degradation. The data with the LCS-HA construct, which is expressed under a constitutive promoter, speak against a GOLPH3 effect on LCS transcription; moreover, we found no differences in endogenous LCS (encoded by *B4GALT5*) mRNA levels under GOLPH3 OE or KD conditions (**Fig. S4A**). The data thus indicate that GOLPH3 OE does not act by enhancing LCS transcription and suggest instead that GOLPH3 OE inhibit LCS degradation.

Based on analogy with the yeast system (Tu et al., 2008), we hypothesized that in GOLPH3-KD cells LCS crosses the Golgi and reaches the lysosomes, and that GOLPH3 OE instead prevents transport to lysosomes. To test this possibility, we monitored the trafficking of LCS through the secretory pathway using LCS-GFP_RUSH and video microscopy. While neither GOLPH3 KD or GOLPH3 OE affected the transport of LCS-GFP_RUSH from the ER to the Golgi (**Fig. 3E and movie S1**), GOLPH3 markedly impacted the transport of LCS-GFP_RUSH through the Golgi. Thus, in control cells, LCS-GFP_RUSH persisted in the Golgi region for ~ 2.5 hours before dissipating into punctate cytosolic structures that could be counter-stained with the lysosomal marker LAMP1 (**Fig. S4B**), and then disappeared; in GOLPH3 KD cells LCS-GFP_RUSH rapidly colonized the LAMP1 positive structures within about 1 hour (with most of this time being spent in the transport between ER and Golgi and through the TGN to the lysosomes); and in GOLPH3-OE cells, LCS-GFP_RUSH remained at the Golgi for up to 10 hours (longest recording time) and never made it to LAMP1 positive puncta (**Fig. 3E, Fig. S4B** and **movie S1**).

These data suggest that, in control cells, a fraction of LCS constantly escapes the Golgi and reaches the lysosomes where it is degraded. This results in a relatively short half-life (roughly 90 min) of this enzyme. GOLPH3 OE strikingly enhances the Golgi retention and residence time of LCS, impeding its arrival at the lysosomes, while GOLPH3 KD greatly decreases the Golgi retention of LCS and speeds up its degradation.

To verify this, we evaluated the LCS-GFP_RUSH protein levels during synchronized transport by western blotting. In control cells the LCS-GFP_RUSH protein levels started to decline after acquiring full glycosylation features, *i.e*., after arrival at the Golgi, with a half-time of ~ 2-3 hours. In GOLPH3-KD cells this decline was faster (half-life < 1 hour), while in GOLPH3-OE cells the decline was extremely slow, or not detectable, for up to 8 hours after release from the ER (longest time point tested **Fig. 3F, G**). Control experiments with the non-GOLPH3 client GalT1 in its GFP_RUSH form showed that this enzyme is not affected by GOLPH3 OE or KD (**Fig. S4C**).

Similar experiments conducted in the presence or absence of the lysosomal degradation inhibitor bafilomycin (Yoshimori et al., 1991), gave results in agreement with the above conclusions. Thus, LCS is retained in the Golgi with an efficiency that depends on the levels of GOLPH3; and once escaped the GOLPH3-dependent retention mechanisms, it reaches the lysosomes and is degraded (**Fig. 3H, I**).

Finally, we examined the effects of GOLPH3 on the other client enzymes, *e.g*., Gb3S, GM3S and GD3S for which we used HA enzyme constructs due to lack of suitable antibodies. The impact of GOLPH3 on these enzymes was very similar to that observed for LCS: GOLPH3 OE increased and GOLPH3 KD decreased their levels (**Fig. 3J, K** and **Fig. S4D**). In contrast, enzymes not bearing the DGB motif (*i.e*., GalT1; sphingomyelin synthase 1 [SMS1]; glucosylceramide synthase [GCS]; LC3S; GM2S and GM1 synthase [GM1S]) were insensitive to manipulations of GOLPH3 levels (**Fig. S4E, F**).

In sum, these data indicate that GOLPH3 drives the recycling of its clients in proportion to its expression levels, and tends to retain them in the Golgi, preventing their escape to the lysosomes and hence their degradation. Crucially, in control cells, the GOLPH3 dependent retention is suboptimal and allows a fraction of the clients to reach the lysosomes, resulting in a constitutively fast turnover of these enzymes. This makes the client enzymes amenable to regulation by GOLPH3. Thus, GOLPH3 OE (a condition found in several tumors) efficiently retains the client enzymes in the *trans*-Golgi, blocking their arrival at, and degradation in, the lysosomes, increasing their levels (**Fig. 3L**). On the contrary, GOLPH3 depletion leads to escape of the client enzymes to the lysosomes, decreasing their levels.

### GOLPH3 reprograms GSL metabolism through its client enzymes

We next asked whether the effect of GOLPH3 on LCS, Gb3S, GM3S and GD3S (as detected using constructs encoding these enzymes) impacts on GSL metabolism. GSL synthesis begins in the ER, with the synthesis of ceramide (Cer) which is then carried either to the TGN by the lipid transfer protein CERT, where it is converted to sphingomyelin (SM) (Hanada et al., 2003), or it moves by vesicular transport to the *cis* Golgi, where it is glycosylated to produce glucosylceramide (GlcCer). GlcCer then reaches the *trans*-Golgi either via the lipid transfer protein FAPP2 or by vesicular transport (D’Angelo et al., 2007; D’Angelo et al., 2013b) and it is further glycosylated to LacCer by the GOLPH3 client LCS. LacCer is a key branchpoint metabolite that feeds into the initiator enzymes of the complex GSL metabolic pathways (*i.e*., Gb3S, GM3S, LC3S and GM2S, for the globo, ganglio, lacto and asialo series, respectively) (D’Angelo et al., 2013a; Merrill, 2011). The GSL metabolic network and the position of the putative GOLPH3 clients in it are schematically represented in **Fig. S1F**.

We first examined the GSL synthetic flux by [^3^H]-sphingosine pulse-chase experiments (see **methods** for details) in HeLa cells and PHFs. In both cell lines, GOLPH3 OE stimulated the production of selected metabolites downstream LCS, *i.e*., LacCer and Gb3 at the expense of those upstream LCS (*i.e*., Cer and GlcCer); Gb3 was the most abundant metabolite, while GM3 was less abundant, as expected from previous reports (D’Angelo et al., 2007), and was barely modified by GOLPH3 OE. GOLPH3 KD had the opposite effects on these metabolites. Lacto and asialo GSLs were not measurable, again in agreement with previous reports in these cell lines (**Fig. 4A**, and **S5A**) (D’Angelo et al., 2007; Halter et al., 2007). Steady-state mass spectrometry and cytofluorimetric measurements (using ShtxB and Cholera toxin B fragment (ChTxB) (Gill and King, 1975; Jacewicz et al., 1986; Russo et al., 2018) evaluate the PM levels of Gb3 and GM1, respectively) were in agreement with the pulse-chase data. GOLPH3 OE increased the levels of Gb3 (though not those of GM1), and reduced the levels of Cer [in particular of the bioactive C16:0 Cer; (Fekry et al., 2018; Kroesen et al., 2001)] (**Fig. 4B-D** and **Fig. S5B**) while GOLPH3 KD had the opposite effects, *i.e.,* it induced an increase in Cer and GlcCer, and a strong decrease in Gb3 levels (**Fig. 4B-D** and **Fig. S5B**). Importantly, the lipidome analysis of GOLPH3-KD and -OE HeLa cells did not show significant changes in other segments of the lipid metabolism (**Fig. S5C**). The effects of GOLPH3 were similar in HeLa cells and PHFs with minor differences (**Fig. 4A, B** and **Fig. S5A, B**) that can be explained by the different expression levels of LCS and Gb3S in the two lines (see legend to scheme in **Fig. S5D**).

**Figure 4.**
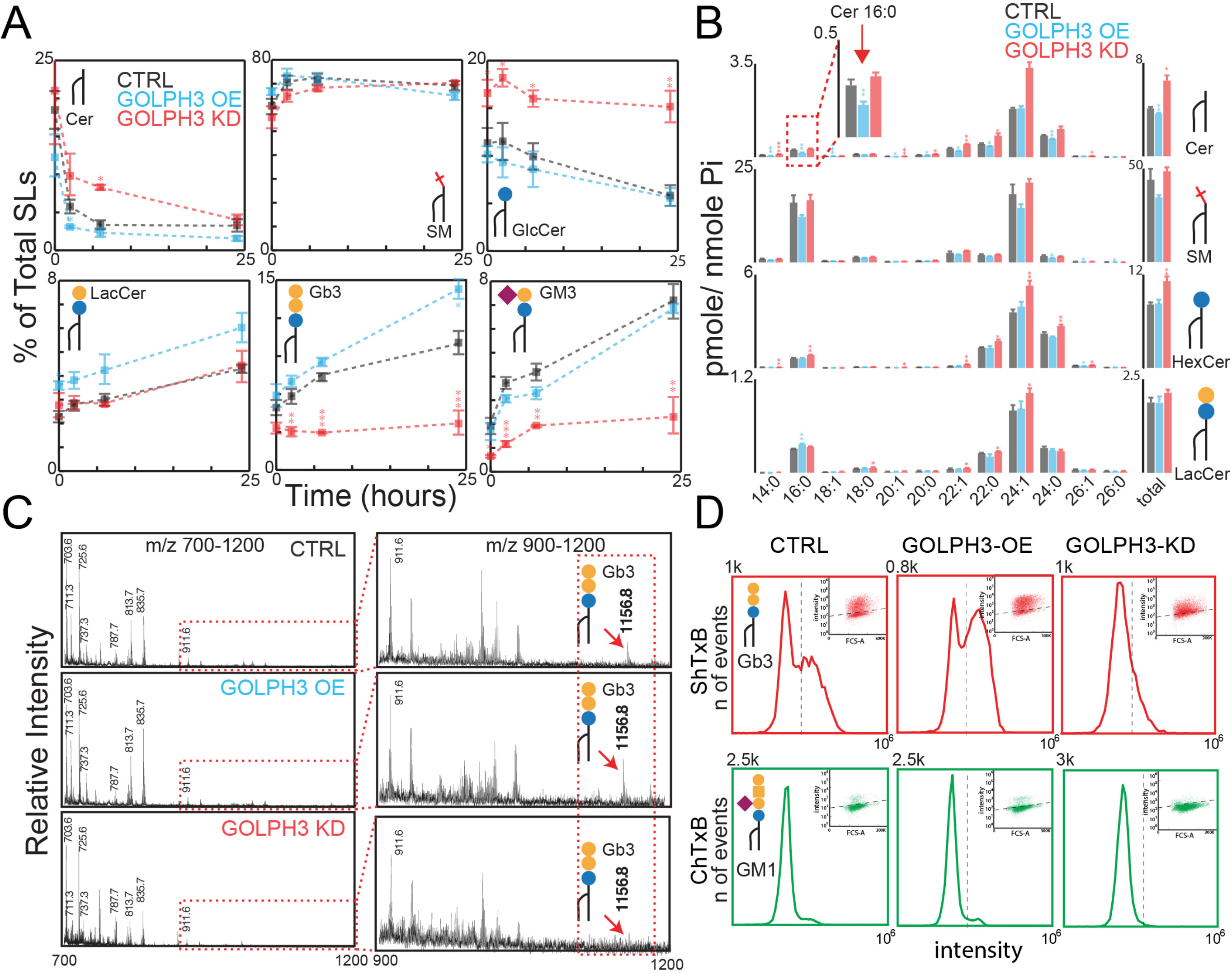
GOLPH3 controls GSL synthesis. (A) Effects of GOLPH3 KD or OE on glycosphingolipid synthetic flux as assessed by [^3^H]-sphingosine pulse (2 hours) and chase (indicated times), lipid extraction and HPTLC separation and radioactive counting (data are means ± SD of at least 3 independent experiments; *p <0.05, **p <0.01, ***p <0.001); (B) Effects of GOLPH3 KD or OE on sphingolipid levels as assessed by LC/MS of HeLa cells (n=3). The dotted red box highlights the effects on C16:0 Cer (data are means ± SD; *p <0.05, **p <0.01, ***p <0.001). (C) Effects of GOLPH3 KD or OE on Gb3 as assessed by MALDI-MS after lipid extraction. Profiles are representative of 3 independent experiments. (D) GOLPH3 KD or OE HeLa cells were processed for ShtxB (upper panels) or ChTxB (lower panels) staining followed by flow cytometry analysis. Flow cytometry distributions, representative of 3 independent experiments, and the relative scatter plots are shown for each condition.

These results can be rationalized by a) examining the position of LCS, Gb3S, GM3S, and GD3S in the GSL metabolic map in **Fig. S5D**; and b) considering the gene expression level of these enzymes in our model cells (see Discussion). Based solely on the position of the GOLPH3 clients in the GSL synthetic pathway one would predict that GOLPH3, by increasing the levels of its clients, should accelerate the production of LacCer and of metabolites of both the globo (Gb3) and ganglio (GM3, GD3) series, at the expense of Cer and of other complex metabolites. However, HeLa cells and PHFs express predominantly globo series synthesizing enzymes, which accounts for the lack of effect on other GSLs series. Thus, our data on GSL metabolism are consistent with the GOLPH3-induced inhibition of lysosomal degradation of its clients as well as with the relative expression of GSL synthetic enzymes. A further notable consequence of these results is that they indicate that GOLPH3 exert similar effects on its clients both as endogenous proteins *(e.g.,* LCS and Gb3S in this case) and as transfected constructs.

Given the key position of the GOLPH3 client LCS in GSL metabolism (see **Fig. S5D**) we examined whether manipulating its levels recapitulates the GSL changes induced by GOLPH3. As shown in **Fig. S5E, F**, LCS OE increased the levels of both LacCer and Gb3, although with a stronger effect on the former and reduced specific Cer species, while LCS KD had the opposite effects. These effects of LCS recapitulate those induced by GOLPH3 in parallel experiments. Moreover, the removal of LCS inhibited the effects of GOLPH3 on LacCer and Gb3. Thus, LCS mimics GOLPH3 with regard to GSL metabolism.

In summary, the main effects of GOLPH3 on GSL metabolism in our model cells are to enhance LacCer and Gb3 levels and to reduce the levels of GlcCer and Cer. These effects were mimicked by LCS. We note that while in our model cells GOLPH3 and LCS have no effects on GSL metabolic branches other than the globo series these are likely to be impacted in cell lines where GSL metabolism is dominated by synthetic enzymes of the ganglio or other complex series.

### LCS and Gb3S mediate the effects of GOLPH3 on mitogenic signaling and tumoral cell growth

The globo-series GSLs produced by the LCS-Gb3S metabolic axis, that is enhanced by GOLPH3 in our model cells, have been reported to activate growth factor receptors and integrin signaling (Chuang et al., 2019; Park et al., 2012; Steelant et al., 2002). These effects resemble those induced by the over expression of GOLPH3, namely, activation of mitogenic pathways (Scott and Chin, 2010; Zeng et al., 2012), including the PI3K, Akt, mTOR and p21^waf^ cascade and of cell growth (Scott et al., 2009; Zeng et al., 2012). We thus examined whether the mitogenic function of GOLPH3 is, at least in part, mediated by its effect on GSL metabolism. Because of the pivotal role of LCS in GSL metabolism, we first compared the effects of LCS OE or KD on these signaling pathways and on cell growth. Remarkably, LCS OE in HeLa cells and PHFs mimicked the stimulatory effects of GOLPH3 OE on Akt and mTOR signaling (*i.e.*, it increased phosphorylation of Akt [at Thr308 and Ser473], and of both p70S6k and 4e-bp1), as well as on the expression of p21^waf^. Conversely, the depletion of LCS mimicked GOLPH3 KD*, i.e.,* depressed the basal activity of the Akt-mTOR pathway and enhanced the expression of p21^waf^ (**Fig. 5A**, and **S6A**). Moreover, LCS recapitulated the effects of GOLPH3 also in assays of cell proliferation as judged by growth in adherence and in soft agar (**Fig. 5B, C**).

**Figure 5.**
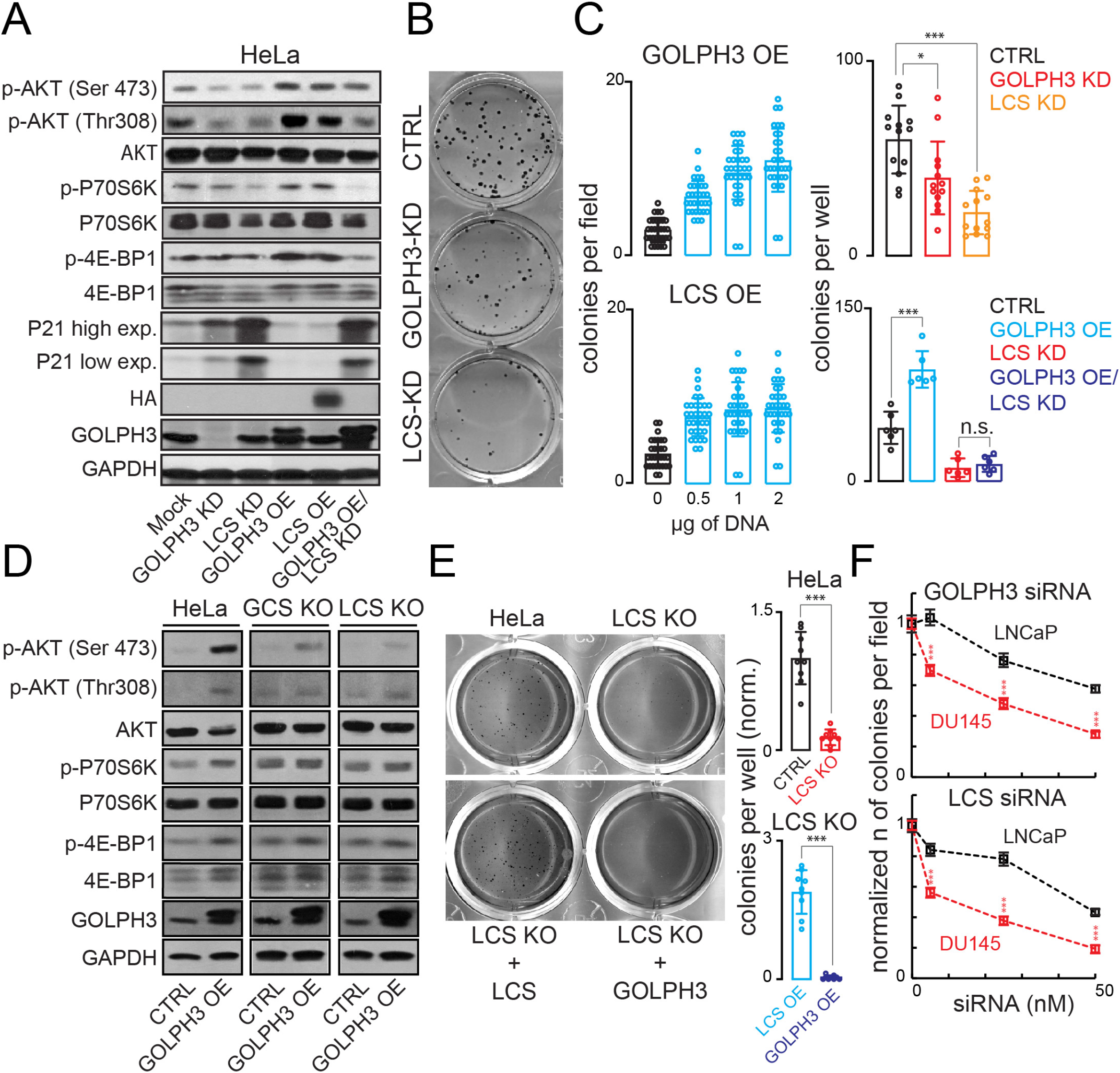
GOLPH3 impacts growth signalling through LCS. (A) Cell lysates from HeLa cells KD or OE for either GOLPH3 or LCS were processed for SDS-PAGE and Western blotting with antibodies aimed at evaluating the activation status of the mTOR pathway. Data are representative of at least 3 independent experiments. (B) Clonogenic assay of HeLa cells in adherence. 250 HeLa cells KD for either GOLPH3 or LCS were allowed to form colonies for 10 days. Afterward, colonies were fixed and stained as detailed in methods. (C) Soft agar colony formation assay of HeLa cells transfected with different concentration of either GOLPH3 (left upper graph) or LCS (left bottom graph) encoding plasmids. Soft agar colony formation assay of HeLa cells KD for either GOLPH3 or LCS (right upper graph). Soft agar colony formation assay of HeLa cells KD for LCS and GOLPH3 OE alone or in combination with LCS KD (right bottom graph). Colonies were stained and quantified as described in methods; (individual data points are shown the graphs; data are means ± SD; *p <0.05, **p <0.01, ***p <0.001, n.s. = not significant). (D) Protein lysates from parental (left), GCS-KO (middle), and LCS-KO (right) HeLa cells, CTRL or GOLPH3-OE were processed for SDS-PAGE and WB with the indicated antibodies. Data are representative of at least 3 independent experiments. (E) Soft agar colony formation assay of parental and LCS-KO HeLa cells expressing LCS-HA or GOLPH3. Colony number was quantified; (individual data points are shown the graphs; data ate means ± SD; ***p <0.001). (F) Quantification of soft agar colony formation assay of LNCaP and DU145 prostatic cancer cell lines treated with increasing concentrations of siRNA targeting GOLPH3 or LCS. (data are means ± SEM; ***p <0.001).

We then asked whether GSLs synthesis is necessary for GOLPH3-induced mitogenic effects. To this end, we examined the effects of GOLPH3 OE in LCS KD as well as in GCS and LCS knock out (KO) HeLa cells (Yamaji and Hanada, 2014). As expected, in both GCS and LCS KO cells the production of complex GSLs was strongly inhibited (**Fig. S6B**). Under these conditions (and in LCS-KD HeLa cells), GOLPH3 OE was unable to induce activation of Akt/mTOR signaling and to stimulate anchorage-independent growth (**Fig. 5A, C-E**). Overexpressing LCS reestablished GSLs production (**Fig. S6B**) and promoted growth in LCS-KO cells; however, it had no effect in GCS-KO cells (not shown) Iindicating that, as expected, LCS requires the presence of the upstream metabolite GlcCer to induce growth, and ruling out potential spurious effects of LCS through reactions on GSL-unrelated substrates. Notably, LCS-KO cells could be induced to grow also by the expression of an unrelated oncogene *i.e*., RAS-Val12 (Sweet et al., 1984) (**Fig. S6C**), indicating that LCS is essential for GOLPH3 dependent growth, but not for growth in general.

Then considering that GOLPH3 enhances both LCS and Gb3S and thus the production of the bioactive GSL Gb3, we asked whether the LCS product LacCer needs to be converted into Gb3 to activate mitogenic signaling. To test this possibility, we depleted Gb3S and monitored the signaling response to GOLPH3 OE. As shown in **Fig. S6D, E**, depletion of Gb3S inhibited the mitogenic signaling and anchorage independent growth induced by GOLPH3 OE. In parallel experiments the depletion of Gb3S inhibited the mitogenic effects of LCS OE (not shown). Thus, the whole GCS-LCS-Gb3S GSL metabolic axis that leads to Gb3 production is needed for the mitogenic effects of GOLPH3 in our model cells. In contrast, the KD of the enzymes initiating the other series (ganglio, lacto and asialo) had minor and non-reproducible effects on GOLPH3-induced mitogenic signaling in line with the barely detectable levels of these GSLs in our model cell lines.

We finally asked whether the GSL-dependent mechanism of action of GOLPH3 in cell proliferation applies to cancer cell lines exhibiting *GOLPH3* amplification. To test this, we first examined two prostatic cancer cell lines, LNCaP and DU145, of which only the latter shows *GOLPH3* amplification [according to the cBioPortal database [http://www.cbioportal.org/; (Gao et al., 2013)]. The DU145 line showed higher levels of both GOLPH3 and LCS as well as of complex GSLs (among which Gb3) compared to LNCaP cells (**Fig. S6F, G**). Moreover, the DU145 line displayed stronger basal phosphorylation of the mTOR substrates p70S6k and p4e-bp1 than LNCaP cells (**Fig. S6F**). All these features are consistent with the genomic amplification and the high GOLPH3 levels in the DU145 line. When we depleted LCS (or GOLPH3) in either the DU145 or in the LNCaP cell lines (**Fig. S6H**), the growth of the DU145 cells was potently inhibited, while LNCaP cells were much less affected, as judged by the soft agar colony formation assays (**Fig. 5F**). Thus, *GOLPH3* amplification and GOLPH3 overexpression correlate with increased LCS protein levels and with the requirement for LCS for growth, in line with the above results obtained in HeLa cells.

### LCS and GOLPH3 levels correlate in human cancers

The above data indicate that the increase in LCS is a key step in the mitogenic effect of GOLPH3 in experimental cells. If the mechanism identified in this study operates in human cancer, a correlation should be detectable between GOLPH3 and LCS expression levels in biopsies from cancer patients. The *GOLPH3* gene has been originally reported to be amplified in different tumor types, with a particularly high frequency (56%) in non-small cell lung cancer (NSCLC) (Scott et al., 2009), and our database search for GOLPH3 copy number variations (CNVs) in large cohorts of lung cancer patients returns (when using stringent defining criteria) significant, though lower, amplification frequencies (~10%) [cBioPortal; http://www.cbioportal.org/; (Gao et al., 2013)] (**Fig. S6I**). In addition, the GOLPH3 protein levels have been reported by many studies to be higher than in healthy lung tissue in > 70% of tumor biopsies from NSCLC patients, and to correlate with poor survival (Tang et al., 2018; Tang et al., 2016). Using a tissue microarray (TMA) prepared from 72 surgical specimens from NSCLC patients (22 females and 50 males with median age at surgery of 63.5 years) (**Fig. 6A**), we examined the correlation between the levels of GOLPH3 and those of LCS. We analyzed GOLPH3 CNVs by fluorescence in situ hybridization (FISH) (**Fig. 6B**) and GOLPH3 and LCS protein levels by immunohistochemistry (IHC) (**Fig. 6C, D**). As shown in **Fig. 6B, C** and in **Table S1**, ~12% and ~40% of the analyzed samples had a *GOLPH3* CN > 6 (amplification) and 4 respectively, and > 60% of the analyzed samples displayed clearly measurable GOLPH3 protein expression levels, in line with previous reports (Tang et al., 2018; Tang et al., 2016). When we examined the LCS protein we found that cancer samples with high GOLPH3 protein levels showed also high LCS protein levels (~65% of cases; **Fig. 6C, D**), while only ~20% of the GOLPH3 weak or negative samples had moderate to high LCS expression levels (**Fig. 6C, D** and in **Table S1**). Overall, we observe a significant (χ^2^ p-val < 0.0001) correlation between GOLPH3 and LCS protein levels, suggesting that the mechanisms by which GOLPH3 controls LCS levels operates in human cancer patients.

**Figure 6.**
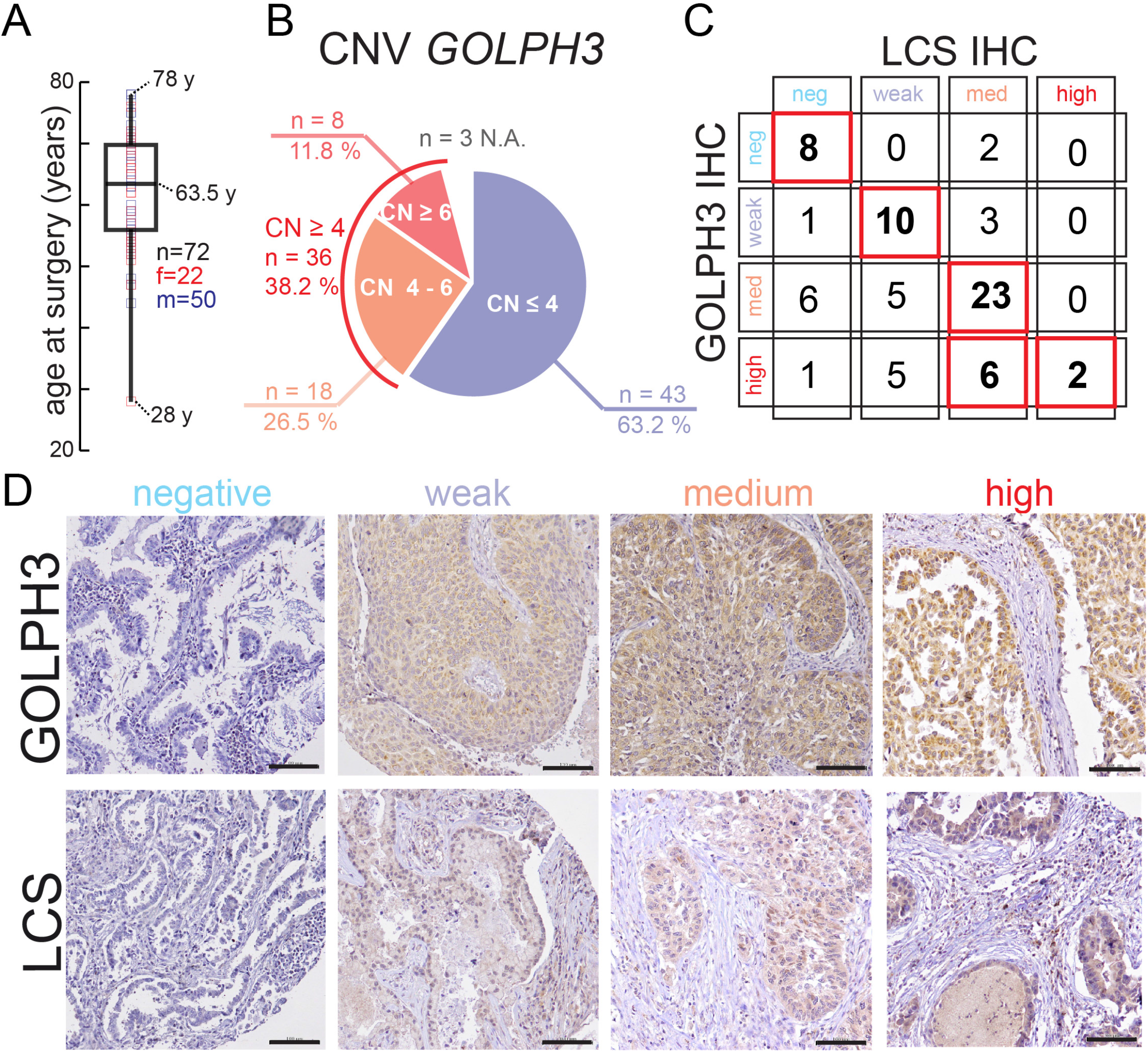
GOLPH3 and LCS protein levels correlate in NSCLC tissue samples. Surgical specimens were collected from 72 non-small cells lung cancer (NSCLC) patients. In (A) age at surgery and sex distribution of the patients are reported. Data are depicted as a box plot. (B) Copy number (CN) variations in *GOLPH3* gene were assessed in cancer tissue section by DNA FISH as described in **Methods** and are expressed as a pie chart. (C) Cancer tissue sections were processed for IHC using antibodies against endogenous GOLPH3 and LCS. The IHC intensity of the staining for each section and for both the tested proteins was classified according to the criteria described in **Methods**. A contingency table for the GOLPH3 and LCS staining is reported. (D) Representative images of NSCLC sections stained for GOLPH3 and LCS. Bar, 100 μm.

## Discussion

We have used GOLPH3, an oncoprotein that operates as an adaptor in the retro-transport of glyco-enzymes across Golgi *cisternae*, as a molecular tool to unravel the links among intra-Golgi transport, glycosylation and cellular responses.

We find that GOLPH3 operates through the following sequence of mechanistic steps. First, GOLPH3 selectively binds a set of glyco-enzymes that includes a functionally coherent group of clients with key roles in the sphingolipid glycosylation pathway. Through this binding GOLPH3 recycles its client enzymes across the Golgi *cisternae* acting as a component of cisternal maturation. The rate of GOLPH3-dependent recycling regulates the levels of the client enzymes by hindering their transport to the lysosomes and hence counteracting their lysosomal degradation. Thus, in cells that overexpress GOLPH3 *e.g*., tumors lines where *GOLPH3* is amplified, the recycling and Golgi retention of GOLPH3 client enzymes becomes very efficient, leading to a marked increase of their levels. Some of the GOLPH3 clients are placed at critical bifurcation points in the GSL metabolic network, so that their increase reprograms the GSL metabolism to decrease growth-suppressing and overproduce growthpromoting metabolites. These lipid metabolic effects are to a large extent responsible for the GOLPH3-driven mitogenic signaling and tumor cell growth and appear to operate also in GOLPH3-dependent human tumors.

Some of these observations carry important implications about the general organization and functional significance of intra-Golgi transport. One is the combination of the findings that GOLPH3 behaves as a component of the maturation mechanism and that it also selectively acts on a functionally coherent enzymatic module. This is inconsistent with the previous cisternal maturation model as a housekeeping process acting homogeneously on all glycoenzymes, and introduces a new view of Golgi maturation by which this process is an assembly of recycling mechanisms acting on different glycosylation pathways with a potential role in the control of cellular glycosylation patterns. Another is the mechanism by which GOLPH3 increases the levels of its client enzymes through a lysosomal degradation-regulated process, which represents a novel pathway for modulating Golgi glycosylation. Some salient mechanistic aspects as well as the physiological and pathological significance of these findings are discussed below.

### GOLPH3 recycles selectively a functionally-coherent glyco-enzyme module yet behaves as an integral component of the cisternal maturation mechanism

A key feature of GOLPH3 is the ability to interact with a selected set of glyco-enzymes with key growth-related roles. This interaction is mediated by a recognition motif in the cytosolic tail of the client enzymes that is similar to the motif recognized by Vps74p in yeast but that contains a second component with one or more positively charged residues upstream of the LxxR sequence (Tu et al., 2008). The cause of this divergence between yeast and mammals is unclear but it might relate to the necessity of variants of the enzyme adaptor recognition motifs to control the vast repertoire of glycosylation pathways emerged during evolution (Gagneux et al., 2015).

Beyond these mechanistic aspects, of broad significance is the concurrence of two findings related to the GOLPH3-enzyme interaction. The first is that through this binding, GOLPH3 controls the activity of a group of enzymes that catalyze sequential reactions at key branchpoints of the GSL metabolic pathways (see map in **Fig. S1F**). By doing so, GOLPH3 acts as a master regulator of GSL metabolism. This represents, to our knowledge, the first case of a Golgi enzyme adaptor that operates on a functionally coherent set of glyco-enzymes to regulate specific cellular functions. A recent study reports that 5 GalnT enzymes involved in the initiation of O-linked glycosylation are retained by recycling in the Golgi via binding to COPI δ and ς subunits (Liu et al., 2018). These enzymes, however do not delineate a functional glycosylation pathway and their relationship with cisternal maturation remains to be defined. The second is that GOLPH3 and its clients operate as components of cisternal maturation. Cisternal maturation has so far been considered a constitutive mechanism that functions merely to maintain the constancy of the enzymatic composition of *cisternae* during intra-Golgi traffic. The observation that GOLPH3 acts only on a specific enzyme set is incompatible with this model and supports instead an alternative view by which, the recycling process underlying maturation is in fact a mosaic of recycling mechanisms acting on different enzyme modules, possibly by virtue of specific adaptors.

The adaptors and recycling mechanisms are most likely regulated: for instance, GOLPH3 is phosphorylated at multiple positions including in its N-terminal portion (McNulty and Annan, 2008; Mertins et al., 2014), *i.e*., the region of contact with COPI, and phosphorylation by DNA-PK has been shown to modulate GOLPH3 function (Farber-Katz et al., 2014). In addition, GOLPH3 is regulated through its recruitment to the Golgi membranes via phosphatidylinositol-4-phosphate [PtdIns(4)*P*] as well as via transcriptional control (Dippold et al., 2009; Wood et al., 2009). Along this line, and of more general significance, several clusters of signaling kinases have been shown to regulate different glycosylation pathways in specific manners (Chia et al., 2012), suggesting that many if not all of the glycosylation modules are densely controlled by signaling mechanisms. In this scenario, the maturation process may orchestrate the glycosylation pathways to change glycosylation patterns according to functional needs, and may therefore also, in case of malfunctions, cause glycosylation imbalances that lead to disease, including cancer. Thus, the altered glycosylation frequently associated with cancer might both result from, and contribute to, the pathological signaling patterns typical of this disease (Chia et al., 2016; Nguyen et al., 2017; Pinho and Reis, 2015; Stanley, 2011; Stowell et al., 2015). Our data suggest that the identification of the recycling mechanisms operating on the various enzyme modules and of the associated regulatory mechanisms will contribute significantly to our understanding of the role of Golgi dynamics in physiology and pathology.

### GOLPH3 increases the levels of its clients by promoting their intra-Golgi recycling and Golgi retention, hence preventing their arrival at the lysosomes

It has been generally thought that the levels of Golgi enzymes are regulated transcriptionally, rather than by the Golgi trafficking, and that the Golgi machinery might affect glycosylation by altering the localization of the enzymes in the stack.

We find instead that GOLPH3 regulates the levels of its clients through a novel lysosomal degradation-based mechanism that depends on the efficiency of the GOLPH3-driven recycling. Degradation-dependent regulatory mechanisms have been shown to control the levels of functionally crucial proteins in several natural processes. Among these, well characterized are the cases of p53 and β-catenin (Jang, 2018; Varshavsky, 2005), both of which are kept constitutively low by targeted ubiquitination and proteasomal degradation, but can be increased rapidly through signal-triggered inhibition of degradation (Jang, 2018; Varshavsky, 2005). The case of the GOLPH3 clients is conceptually similar but relies on a different mechanism that can be rationalized in the context of cisternal maturation. In the maturation scenario, the localization of GOLPH3 clients in the stack depends on two opposing but normally well-balanced processes, namely, the anterograde progression of *cisternae* on the one hand, and the GOLPH3-driven retrotransport of its client enzymes on the other. The former tends to carry these enzymes through the Golgi and to the lysosomes, and the latter to retain the GOLPH3 clients in the Golgi, shielding them from degradation. When GOLPH3 is overexpressed (a situation found in numerous cancers (Scott et al., 2009), the balance between the two above processes is altered. Thus, under high GOLPH3 expression regimens, the client enzymes are recycled more efficiently, and do not reach the lysosomes, as a consequence they are degraded less, and their levels increase markedly. Conversely, when GOLPH3 is depleted, they are degraded rapidly. Thus, GOLPH3 acts as a rheostat that controls the flow of its clients to the lysosomes, thereby regulating their levels.

Remarkably, in control cells, GOLPH3 driven recycling rates appear to be suboptimal and to allow a fraction of the GOLPH3 clients to continuously leave the stack for the lysosomes, where they are degraded. This results, at least for the GOLPH3 client LCS, in a half-life of ≈ 2 hours (shorter than for most cellular proteins). This relatively rapid turnover creates the conditions for GOLPH3 OE to increase the levels of its clients, through the above-described degradation-based mechanism. Importantly, this high basal turnover of GOLPH3 clients is not a peculiarity of HeLa cells. Data consistent with this degradation-based mechanism have been obtained also in PHFs and are in line with observations in prostate cancer cell lines and in cancers from human patients. The suboptimal rate of enzyme recycling and of Golgi retention efficiency appears, therefore, to be a programmed feature of the GOLPH3 recycling machinery designed to render the system amenable to regulation by GOLPH3.

To our knowledge, this is the first example of a mechanism by which Golgi dynamics control the levels of glyco-enzymes and thereby regulate cell growth. Changes in glycosylation patterns are very frequent in cancer cells. They have been so far interpreted as being due to a changed glyco-enzyme gene expression (Dall’Olio and Trinchera, 2017). Regulated-degradation processes, like the one described here, should now be considered as a possible alternative mechanism.

### GOLPH3 reprograms GSL metabolism to promote mitogenic signaling and cell proliferation

A few hypotheses have been put forward to account for the oncogenic action of GOLPH3 (Farber-Katz et al., 2014; Halberg et al., 2016; Isaji et al., 2014; Rizzo et al., 2017; Scott and Chin, 2010). Our data indicate that the effect of GOLPH3 on GSL metabolism represent a central component of the GOLPH3 pro-growth effects, but do not exclude a role of additional mechanisms in the action of this adaptor.

Among the GOLPH3 clients, LCS, Gb3S, GM3S and GD3S are likely to play the main role in growth induction as they catalyze reactions in GSL metabolism that can lead to the production of growth-promoting lipids. Mapping these GOLPH3 clients on the GSL metabolic network (**Fig. S1F, S5D**) shows that these enzymes sit at crucial metabolic branchpoints, with predictable stimulatory effects on growth. Thus, LCS channels the metabolic flux towards Gb3S and GM3S, the initiator enzymes of the globo and ganglio series, respectively, at the expense of Cer, a tumor suppressor and growth inhibitor (Hannun and Obeid, 2008; Reynolds et al., 2004). The increase in Gb3S initiates the synthesis of globo series GSLs, some of which are known to activate integrins and growth factor receptors (Steelant et al., 2002), while GM3S initiates the synthesis of gangliosides, and GD3S converts the growth-inhibiting GM3 (Bremer et al., 1986; Coskun et al., 2011) into the growth promoting GD3 (Ohkawa et al., 2010). By acting on these enzymes, GOLPH3 OE can reprogram the GSL pathway towards a pro-growth configuration.

Importantly, however, the specific GSLs configuration induced by GOLPH3 OE in each cell line depends on the transcriptional control of the levels of the GSL enzymes, which varies across cell types, tissues, and tumor types. For instance, in the neural tissue, gangliosides are prevalent, so the effects of GOLPH3 are likely to be expressed through increases in the levels of GD3, a characterized activator of growth receptors, and are thus likely to induce growth via this pathway (Ohkawa et al., 2010). In our cell models, the globo pathway is prevalent; and we indeed find that GOLPH3 induces growth through effects on the globo series GSLs. Although the precise mechanism by which they exert their effects remains to be determined, Gb3 and the downstream metabolites Gb4, and sialo-Gb5 have been shown to cluster in GSL rich domains at focal adhesions (Steelant et al., 2002), where they bind to and activate integrins and RTK receptors to promote PI3K-Akt-mTOR signaling (Chuang et al., 2019; Furukawa et al., 2019; Hakomori Si, 2002; Park et al., 2012; Steelant et al., 2002; Wegner et al., 2018). An additional contributing factor to the pro-growth effects of GOLPH3 that is independent of GSL pathway being affected, is the decrease of Cer, a characterized tumor suppressor and inhibitor of cell proliferation (Hannun and Obeid, 2008; Reynolds et al., 2004).

Although the effects on GSL metabolism are a major component of the pro-growth action of GOLPH3, the are unlikely to fully explain the oncogenic action of this adaptor. Other GOLPH3 clients might contribute to, or be necessary for, these effects of GOLPH3. For instance, ST6GAL1, a GOLPH3 client and a promiscuous *trans* Golgi capping enzyme acting on both GSLs and proteins has been proposed to mediate the effect of GOLPH3 on cell migration through sialylation of integrins (Isaji et al., 2014). Moreover, the role of a few potentially relevant putative GOLPH3 clients involved in the synthesis of proteoglycans, or of the Lewis antigens, which is involved in cell extravasation and invasion (Hakomori, 1989; Hakomori and Zhang, 1997), or in N- or O-linked glycosylation, remains to be examined. Finally, also enzyme-unrelated activities of GOLPH3 have been proposed to contribute to its oncogenicity. Among these, the GOLPH3 effects on cargo export from the TGN, and on the DNA damage response have been thoroughly characterized (Dippold et al., 2009; Farber-Katz et al., 2014; Halberg et al., 2016; Rahajeng et al., 2019; Xie et al., 2018). It will be interesting to examine how these mechanisms synergize with the effects of GOLPH3 on GSL metabolism described in this study.

## Perspectives

Altogether, this study presents the first example of a mechanism by which intra-Golgi transport regulates glycan assembly and associated biological functions. The property of GOLPH3 to control selectively an enzymatic module, the potentiation of its clients over competing enzymes by a degradation-regulated mechanism, and hence the production of certain glycans over others, represents a new paradigm in the control of Golgi glycosylation. The current findings might thus provide a roadmap for the analysis of the role of the Golgi in glycosylation based on the identification and analysis of the Golgi enzyme adaptors and of the biosynthetic modules they control. In addition, they introduce a new view of cisternal maturation that implies a potential role of this mechanism as a regulator of glycosylation, with important potential consequences in physiology and pathology.

On a more translational ground, the clinical data suggest that the mechanisms described here operate in tumors of human patients. GOLPH3 and its upstream regulators (*i.e.*, the *trans* Golgi located PtdIns(4)*P* signaling circuit) are emerging as clinically relevant entities in a number of frequent and poorly treatable human cancers (Halberg et al., 2016; Scott et al., 2009). In this study we provide data on what is most likely the prevalent oncogenic mechanism of GOLPH3 (and possibly of oncogenes that rely on GOLPH3 for their activity), which is bound to have significant medical and pharmacological relevance.

## Supporting information

Sipplementary Figures

## Acknowledgments

We thank Antonella De Matteis, Francesca Carlomagno, Lina Obeid for valuable discussions. We thank Nina Dathan for help with cloning and construct preparation. We thank Francesco Russo for Bioinformatic support. We thank Pasquale Barba for assistance and the FACS Facility of the Institute of Genetics and Biophysics (IGB-CNR, Naples). We thank Gabriele Turacchio for EM technical assistance and the Bioimaging Facility of the Institute of Protein Biochemistry. We thank Florence Pojer, Kelvin Lau and the Protein production and structure core facility – PTPSP at EPFL for technical support on protein purification and ITC experiments. We thank Stuart Kornfeld and Balraj Doray for the sucrose isomaltase encoding plasmid. R.R. acknowledges financial support from Fondazione Italiana per la Ricerca sul Cancro (FIRC Fellowship 15111). A.L. acknowledges financial support from the Italian Cystic Fibrosis Research Foundation (Project #6), AIRC (Projects IG 15767 and IG 20786), the Italian Node of Euro-Bioimaging (Preparatory Phase II – INFRADEV), TERABIO the MIUR PON Project IMPARA and the POR Capania projects 2014-2020 C.I.R.O. and S.A.T.I.N. G.D.A. acknowledges financial support from EPFL institutional fund, Kristian Gerhard Jebsen Foundation and from the Swiss National Science Foundation (SNSF) (grant number, 310030_184926).

## Authors contributions

RR developed the idea, conducted most of the experiments and contributed to write the manuscript; DR participated in the development of the idea and conducted the experiments on signalling and peptide binding; KK conducted the experiments with yeast cells; PS, Conducted the GalT1-RUSH experiment and contributed to cell signaling and proliferation experiments; BL was involved in the setup of the RUSH synchronization protocol of LCS-GFP_RUSH and EM assessment of GOLPH3 distribution; DS assisted RR in conducting the experiments in an initial phase of the project; MZ conducted bioinformatics analysis of Golgi enzymes tails; AV Conducted the ITC experiments; PP and VK, assisted RR in conducting the experiments the final phase of the project; LC conducted MALDI-MS and lipidomics measurements; GB and FP generated RUSH constructs, FZM, GA, and CV conducted in TMA experiment; PH Generated biotinylated peptides; LM expressed and purified recombinant GOLPH3; DM, assisted RR in conducting the experiments in an initial phase of the project, GB and AB analysed and interpreted the TMA data; HC and UM generated and provided LCS antibody and analyzed glycosylation data; TY and KH generated and provided the GCS and LCS KO HeLa cells; SP, contributed to conception and design, analysis and interpretation of data and contributed to write the manuscript; YAH helped with analysis and interpretation of lipids MS data and contributed to write the manuscript; AN was Involved in data analysis and interpretation of yeast experiments; DC contributed to development of the idea and to the critical analysis of the data throughout the entire project; GDA and AL developed the idea, designed and supervised the entire project, and wrote the manuscript.

