## Supplementary material for "The Glyco-enzyme adaptor GOLPH3 Links Intra-Golgi Transport Dynamics to Glycosylation Patterns and Cell Proliferation"

Supplementary Figure Legends

### Figure S1. GOLPH3 binds a compound motif in the cytosolic tail of specific GSL synthetic enzymes

(A) Simplified scheme of the cisternal progression maturation process in a mammalian cell. Cisternal progression maturation can be seen as the integration of two processes, cisternal progression and cisternal maturation. We note that while the core of the overall process, namely, the fact that *cis cisternae* form, mature into *trans* elements, and then finally disassemble, is conserved across eukaryotes, the morphology and organization of various steps can vary across cell types. In brief, cargo proteins (black dots) reach the *cis* Golgi from the ER within membranous carriers which then fuse into the ER-Golgi Intermediate Compartment (red) and form new *cis cisternae* (blue). The new cisternae compositionally convert into medial and *trans* cisternal elements through the backward recycling (via COPI coated vesicles) (grey arrow) of glyco-enzymes (red, green and grey dots) and other resident proteins from distal *cisternae*, and then finally disassemble into carriers destined to the PM or other destinations (cisternal progression). The recycling of glyco-enzymes occurs in synchrony with cisternal progression to maintaining the compositional homeostasis of the stack during cisternal progression (cisternal maturation). The most significant morphological variants of this process are as follows: mammalian cells have the ERGIC between ER and Golgi while yeast cells do not; *cisternae* are organized in stacks in most cells while they exist as separate entities moving through the cytosol in yeast; the mode of assembly and disassembly of cisternae at the *cis* and *trans* Golgi face, respectively, differs across mammalian and yeast cell types. For instance, the TGN detaches in block from the stack in *Pichia pastoris* yeast, while it disassembles gradually into pleomorphic carriers in mammals; in *Pichia pastoris* the trans cisterna appears to mature into the TGN, while in mammalian cells the trans cisterna appears to deliver cargo through anterograde carriers to the TGN. The morphology and dynamics of the formation of new *cis* cisternae may also vary between yeast, mammals and plants. The molecular mechanism underlying cisternal progression maturation is partially understood. Several potential players have been identified. These include components involved in membrane budding, fission, tethering docking fusion as well as proteins that might be specialized for intra-Golgi traffic such as the COGs, golgins and others. Among these proteins, only the COPI coat has been ascribed a well-defined function in cisternal maturation. (B) Glyco-enzymes bearing the LxxR motif in the cytosolic tail grouped according to the pathway in which they are involved. Red, GSLs; Cyan, proteoglycan; Purple, N-glycan; Green O-glycan. (C) Flow cytometry-based lectin screening of GOLPH3-OE HeLa cells. AAL, *Aleuria aurantia* Lectin; SNA, *Sambucus nigra* Lectin; PNA, *Peanut Agglutinin*; RCA-I *Ricinus communis* Agglutinin I; PHA-L *Phaseolus vulgaris* Leucoagglutinin; ShTxB, Shiga toxin 1a B fragment. The glycan specificity of each lectin/ toxin is reported. (D) Hela cells expressing LCS-HA were lysed and LCS-HA was immunoprecipitated by anti-HA antibody. Coimmunprecipitated GOLPH3 was detected in the immunoprecipitated fraction. F.T, flow through fraction; I.P. immunoprecipitated fraction. (E) List of 22 glyco-enzymes bearing the LxxR motif (red) and at least a positively charged residue (cyan) in the 3 to 6 amino acids upstream the LxxR in their cytosolic tail. The coloured squares indicate the metabolic pathways each enzyme is involved in: Red, GSL metabolism; Cyan, glycosaminoglycan assembly; Purple, N-glycosylation; Green O-glycosylation. (F) Schematic representation of GSLs synthesis and of the enzymes (in red) that bear the DGB motif. Note that LacCer is the common precursor for the synthesis of globo, ganglio, lacto and asialo GSL series. (G) Pull-down experiments involving biotinylated peptides of glyco-enzyme cytosolic tails from purified His-tagged GOLPH3 protein. The N-terminal cytosolic tail of the involved glyco-enzymes is shown on top of the blot (LxxR motif in red, positively charged residue in cyan, biotin at the C-terminus). Immunoblotting shows affinity-captured recombinant GOLPH3. (H) Sequences of all the glyco-enzymes cytosolic tails tested for GOLPH3 interaction in this or in other (asterisks) studies. The red box indicates the LxxR motif in red, the cyan box indicates the positively charged residue region.

### Figure S2. Vps74p controls the sub-Golgi distribution and Golgi retention of Mnn9p

(A) GOLPH3-KD HeLa cells were processed for cryoimmunolabeling. Anti-GalT1 antibody (10 nm gold particles) and anti-GM130 antibody (15 nm gold particles, arrowheads) are indicated as markers of the cis side of the Golgi stack. Quantification of the distribution (LD normalised to the *trans cisternae*) of GalT1 across the stack and peri-Golgi vesicles is shown in the graphs (right). Data are means ± SEM. Bar 150 nm. (B) Images of wild-type (left) and *vps74Δ* (middle) yeast cells expressing Mnn9-mCherry (red, Vps74p client) and Sys1-GFP (green, *trans*-Golgi marker). Arrowheads indicate *cisternae* with Mnn9-mCherry and Sys1-GFP. The fraction of positive *cisternae* (Sys1-GFP, Mnn9-mCherry, or both of them) in wild-type and *vps74Δ* yeast cells was quantified and is shown in the right box plots; (n is indicated in the graph; data are means ± SEM; *p <0.05, ***p <0.001). Bar, 1µm. (C) Wild-type and *vps74Δ* yeast cells expressing Mnn9-mCherry (red, Vps74p client) and Sys1-GFP (green, *trans*-Golgi marker) were treated with 500 µg/ml cycloheximide for 60 min and then observed. Images of Mnn9-mCherry and Sys1-GFP in WT (left two panels) and *vps74Δ* (right two panels) yeast cells are shown. (D) Images of wild-type (upper) and *vps74Δ* (lower) yeast cells expressing Mnn9-mCherry (red, Vps74p client) and Sys1-GFP (green, *trans*-Golgi marker) were observed. Representative images are shown on the left. Bars, 1 µm. Middle panels show time-lapse images of selected *cisternae* (boxed areas in the left panels) in wild type (upper) and *vps74Δ* (lower) yeast cells. Right graphs show relative fluorescence intensity changes of Mnn9-mCherry (red) and Sys1-GFP (green) of 11 selected cisternae in wild-type (upper) and 10 selected cisternae in *vps74Δ* (lower) yeast cells. Thick lines in the graphs show representative fluorescence intensity changes. (E) Wild-type (upper) and *vps74Δ* (lower) yeast cells expressing Mnn9-sfGFP (green, Vps74p client) and Vph1-mCherry (red, vacuole marker) were observed. Representative images before photo-bleach are shown on the left. Bars, 1 µm. Images of selected cells before (pre-bleach) and after (0 min and 5 min) photobleaching

Mnn9-sfGFP are shown on the right. Upper panels show merged images of Mnn9-sfGFP and Vph1-mCherry. Lower panels indicate Mnn9-sfGFP localization. Fluorescence signals of Mnn9-sfGFP after photobleaching are emphasized. Outer dotted circles indicate the edges of cells. Inner dotted lines indicate vacuolar regions. (F) Fluorescence recovery after photobleaching (FRAP) of Mnn9-sfGFP fluorescence intensities in 15 wild-type (blue) and 15 *vps74Δ* (red) yeast cells were plotted at every 2 min during observation time in the upper graph. Average fluorescence intensities (thick lines) ± SEM (thin lines) are shown. Fluorescence intensities of Mnn9-sfGFP and Vph1-sfGFP in selected yeast cells were quantified at indicated time after photobleaching. Graph of Mnn9-sfGFP (bottom left) and Vph1-sfGFP (bottom right) fluorescence intensities in wild-type (blue) and *vps74Δ* (red) yeast cells are shown (n is indicated in the graph, data are means ± SEM). (G) The panels show representative images of Vph1-sfGFP in wt (left) and *vps74Δ* (right) yeast cells before (pre-bleach) and after (0min and 5 min) photobleaching. Fluorescence signals of Vph1-sfGFP after photobleaching are emphasized.

### Figure S3. GOLPH3 controls LCS sub-Golgi distribution

(A) GOLPH3-OE (top pictures) or KD (bottom pictures) LCS-HA expressing HeLa cells were fixed and processed for cryoimmunolabeling with an anti-HA antibody (15 nm gold particles) and anti-GOLPH3 antibody (10-nm gold particles, arrowheads) or anti-GM130 antibody (10-nm gold particles, arrowheads) as indicated. Bar, 250 nm. Quantification of the distribution (LD normalised to the *trans cisternae*) of LCS across the stack and in peri-Golgi vesicles is shown in the graphs on the right; data are means ± SEM. Bar, 150 nm. (B) LCS-HA expressing HeLa cells were fixed and processed for cryoimmunolabeling with an anti-GM130 antibody (15 nm gold particles) and anti-GOLPH3 antibody (10 nm gold particles, black arrowheads), or an anti-HA antibody (15 nm gold particles) and anti-GOLPH3 antibody (5 nm gold particles, black arrowheads). Bar, 150 nm. Images are representative of at least 3 independent experiments.

### Figure S4. GOLPH3 controls the localization and levels of selected GSL synthetic enzymes

(A) mRNA levels of *B4GALT5* (LCS) were evaluated in GOLPH3-OE or KD HeLa cells or PHFs by qRT-PCR. (n = 3; data are means ± SD). (B) LCS-GFP_RUSH expressing GOLPH3-OE or KD HeLa cells were fixed at the indicated time points after traffic synchronization and processed for IF: LCS (green), LAMP1 (red); (n = 3; Bar, 20 µm). (C) GalT1-GFP_RUSH expressing GOLPH3-OE or KD HeLa cells were lysed at the indicated time points after traffic synchronization. Protein lysates were processed for SDS-PAGE and immunoblotted with the indicated antibodies (data are representative of three independent experiments). (D) GOLPH3-OE or KD HeLa cells expressing the indicated glyco-enzymes were lysed and processed for WB; data are representative of three independent experiments. (E) GOLPH3-KD HeLa cells expressing the indicated glyco-enzymes were treated with bafilomycin A1 (BafA1, 10 nM) for 16 hr, fixed, and processed for IF: HA (green), LAMP1 (red), and TGN (blue); images are representative of at least three independent experiments. Bar, 20 µm. (F) GOLPH3-KD HeLa cells expressing the indicated glyco-enzymes were fixed and processed for IF. HA (green), GOLPH3 (red) and TGN46 (blue). Bar, 20 µm. Images are representative of at least 3 independent experiments.

### Figure S5. Effects of GOLPH3 and LCS manipulations on lipid metabolism

(A) Effects of GOLPH3 KD or OE on GSL metabolism as assessed by [$^{3}$H]-sphingosine pulse (2 hours) and chase (indicated times), lipid extraction and HPTLC separation and radioactive counting of PHFs (n = 3; data are means ± SD). (B) Effects of GOLPH3 KD or OE on sphingolipid levels as assessed by LC/MS of PHF cells (n = 3; data are means ± SD; *p <0.05, **p <0.01, ***p <0.001). The dotted red box highlights the effects on C16:0 Cer. (C) Phospho-lipidomics of GOLPH3 OE and KD HeLa cells. Data are shown in a mirror-image volcano plot. Individual lipids are color-coded according to the class they belong to. The dotted lines indicate the significance thresholds (i.e., p<0.05 and log2 fold change <-1 or >1). (D) Simplified scheme of the GSL metabolic pathways in which the positions of the verified GOLPH3 clients (LCS, GB3S, GM3S and GD3S) (red) and the effects of GOLPH3 on these enzymatic activities are represented (red arrows). GOLPH3 increases LCS (bold red) and accelerates the conversion of GlcCer to LacCer (thick red arrow), resulting in the increase of LacCer (bold red) and in the decrease of the metabolites upstream LCS, i.e., GlcCer and Cer (blue). GOLPH3 also increases Gb3S and accelerates the conversion of LacCer into Gb3 (thick red arrow). Notably, LacCer may increase or decrease depending on the balance between the enhanced activities of LCS and Gb3S. What remains constant is the GOLPH3-dependent increase of the pool of metabolites downstream LCS (LacCer plus Gb3) and the decrease of the upstream metabolites the Cer/GlcCer pool. GM3S and GD3S are represented (red) in the scheme but they do not play a significant role in the model cells used in this study because the relevant enzymes are minimally expressed. Lacto and asyalo series (grey) do not play significant roles, for the same reason. (E) Effects of LCS KD or OE on sphingolipid levels as assessed by LC/MS of HeLa cells. The magnified box highlights the effects on C16:0 Cer (n = 3; data are means ± SD; *p <0.05, **p <0.01, ***p <0.001). (F) LCS OE HeLa cells processed for ShtxB staining followed by flow cytometry analysis. Representative flow cytometry distributions and the relative scatter plots are shown for each condition. Data are representative of three independent experiments.

### Figure S6. GOLPH3 impacts growth signalling through changing GSL metabolism

(A) Cell lysates from PHFs KD or OE either GOLPH3 or LCS were processed for SDS-PAGE and Western blotting with antibodies aimed at evaluating the activation status of the mTOR pathway and p21. (B) Assessment of GSL synthesis in parental HeLa, GCS-KO, LCS-KO and LCS-KO cells overexpressing LCS, [$^{3}$H]-sphingosine pulse (2 hours) and chase (24 hours), lipid extraction, HPTLC separation, and radioactive counting (n = 3, data are means ± SD). (C) Quantification of soft agar colony formation assay of HeLa LCS-KO cells overexpressing RasVal12 (n is indicated in the graph; data are means ± SEM; ***p <0.001). (D) Lysates from GOLPH3-OE HeLa Gb3S-KD were processed for SDS-PAGE and Western blotting and the levels of p-Akt (Ser473 and Thr308) were evaluated. (E) Soft agar colony formation assay of the experiment in (D) (n is indicated in the graph; data are means ± SEM; ***p <0.001, n.s. = not significant). (F) Cell lysates from prostate cancer cell line, LNCaP and DU145 respectively, were processed for SDS-PAGE and Western blotting with antibodies aimed at evaluating the levels of expression of GOLPH3, LCS and the activation status of the mTOR substrates. (G) Assessment of GSL synthesis in LNCaP and DU145 cell line by [$^3$H]-sphingosine pulse (2 hours) and chase (24 hours), lipid extraction and HPTLC separation and radioactive counting; (n = 2, data are means ± SEM). (H) mRNA levels of both *GOLPH3* and *B4GALT5* (LCS) were evaluated in LNCaP and DU145 cells under control conditions and upon GOLPH3 and LCS downregulation respectively (using 25 nM of siRNAs, **see Fig. 5F**), by qRT-PCR; (n = 3; data are means ± SD). (I) The cBioportal database (http://www.cbioportal.org) for cancer genomics was interrogated for GOLPH3 gene in 32 cancer sample collections corresponding to 26731 fully sequenced cancer samples. The graph represents the alteration frequency of GOLPH3 in each sample collection. Boxes highlight the expression levels of GOLPH3 in Lung Squamous and Adenocarcinoma according to *GOLPH3* ploidy.

**Movie S1: LCS-GFP_RUSH synchronized transport in GOLPH3-OE or-KD HeLa cells**

GOLPH3-OE or KD HeLa cells were transfected to express LCS-GFP_RUSH. After 24 hours of expression, at time 0, DMEM containing cycloheximide (10 mg/mL) and biotin (40 nM) was added to cells. The trafficking of LCS-GFP_RUSH was monitored using confocal microscope. Images were acquired at 5 minutes intervals.

### Star Methods

Contact for Reagents and Resource Sharing

Further information and requests for resources and reagents should be directed to and will be fulfilled by the Lead Contacts: Alberto Luini and Giovanni D'Angelo.

Cell lines and culture conditions.

HeLa-M (human cervical cancer cells, female origin) were obtained from the ATCC. DU145 (prostate cancer cells, human male origin) were a kind gift from Carmen Valente from the Institute of Biochemistry and Cell Biology, CNR, Naples. LNCaP cells (prostate cancer cells, human male origin) were a kind gift from Alfredo Budillon from the Istituto Nazionale Tumori "Fondazione Pascale". HeLa-M, DU145 and LNCaP cells were cultured in RPMI-1640 medium containing 4.5 g/L glucose, 2 mM L-glutamine, 10% fetal calf serum (FCS), and 100 U/mL penicillin and streptomycin. Primary human fibroblasts (PHFs) (normal cells, male origin) were obtained from the Telethon Biobank. The HeLa-mCAT#8 (including TALEN UGCG-KO, and LCS-KO) cell lines (human cervical cancer cells, female origin) were kind gifts from Kentaro Hanada. Primary human fibroblasts, HeLa-mCAT#8, UGCG-KO, and LCS-KO were cultured in DMEM containing 4.5 g/L glucose, 2 mM L-glutamine, 10% FCS, and 100 U/mL penicillin and streptomycin. Cells were grown under controlled temperature and atmosphere.

METHOD DETAILS

*Plasmids and siRNA transfection:*

HeLa cells and PHFs were transfected with Mirus or Lipofectamine® LTX transfection reagents following the manufacturer's instructions (1% FCS in HF medium). Oligofectamine was used for siRNA transfection and performed according to manufacturer's instructions (used at concentrations between 5 and 50 nM for the target gene). Each individual siRNA in the pool was also tested in localization and signaling experiments. The list of siRNA sequences can be found in Table S2.

*Biotinylated peptides:*

The C-terminally biotinylated peptides corresponding to the tails of Golgi enzymes, were synthetized by Dr. Petra Henkein, Charité - Universitätsmedizin Berlin. Their list and sequences can be found in Table S3.

*Generation of constructs and mutants:*

The GOLPH3 siRNA-resistant construct was generated by site-directed mutagenesis. The oligonucleotide primers used for the mutagenesis reaction were: F5′-gggcctcaaggaccgagagggatacacttcattttggaatgac-3′ and R5′-gtcattccaaaatgaagtgtatccctctcggtccttgaggccc-3′. The oligos were ordered for creating siRNA-1# resistant GOLPH3 by changing the nucleotide sequence, but not the codon sequence. The primers were chosen with melting points above 78 °C, and were self-complementary and PAGE-purified. The GOLPH3 cDNA was cloned in pCMV6-XL5 vector (Origene). Pfu Turbo DNA polymerase was used to insert and amplify the 3-point-mutagenized GOLPH3 sequence, amplifying for 18 cycles at 56 °C and extending the mutagenized DNA 16 min at 68 °C at each cycle. The parent strand was digested with DpnI treatment for 60 min at 37 °C. The final product was precipitated, resuspended in 10 mM Tris, pH 8.0, used to transform TOP10 competent cells, and selected on LB + Amp agar. Minipreps were prepared (QIAprep spin miniprep kit, Qiagen) and the sequence was confirmed with internally designed primers: GF: 5′-atgacctcgctgacccag-3′ and GR: 5′-cttggtgaacgccgccac-3′. The GOLPH3 binding pocket single point mutant R90L was generated by site-directed mutagenesis. The oligonucleotide primers used for the mutagenesis reaction were: R90LF5′-ctgtatatcatctggattacttggctgtatgttaattgaattagc-3′ and R90LR:5′-gctaattcaattaacatacagccaagtaatccagatgatatacag-3′. The oligos were ordered for creating a single point mutant in the binding pocket of GOLPH3. The primers were chosen with melting points above 78 °C, and were self-complementary and PAGE-purified. Mutagenesis, minipreps, and sequencing were performed as described above.

The B4GALT5-Myc (a kind gift from Antonella De Matteis, TIGEM, Naples) was used as a template to generate B4GALT5-HA (LCS-HA) cDNA using F5′-gcgcgcgaattcatgcgcgcccgccg-3′ and R5′-cgcgcgctcgag ttaagcgtaatctggaacatcgtatgggtagtactcgttcacctgagccag-3′ primers. Oligos forward and reverse (incorporating a 3′ HA tag) were used for amplifying 50 ng *B4GALT5* cDNA with 1U Phusion polymerase (NEB) in GC Buffer for 30 cycles on an MJ REsearch Peltier Thermal Cycler, annealing at 70 °C to maintain specificity. The PCR fragment was cleaned using the QIAquick PCR purification kit (Qiagen) and cut with EcoRI/XhoI, re-cleaned with the QIAquick PCR purification kit, ligated into EcoRI/XhoI-cut pcDNA4b for 2 hr at room temperature, and plated on LB + amp agar. Minipreps were prepared (QIAprep spin miniprep kit) and sequence confirmed using T7 and BGH rev primers by PRIMM.

*RNA extraction and real-time qPCR:*

After treatment, total RNA was isolated from cells using RNeasy Mini kits (Qiagen) following the manufacturer's instructions. The quality and quantity of RNA was determined by NanoDrop 2000c spectrophotometer (Thermo Scientific) and TAE agarose gel electrophoresis, respectively. Total RNA was reverse-transcribed to cDNA using the QuantiTect Reverse Transcription Kit (Qiagen) and subjected to real-time qPCR using specific primers for the target genes of interest in the presence of LightCycler® 480 SYBR Green I Master Mix (Roche) on a LightCycler® 480 II detection system (Roche). Hypoxanthine phosphoribosyltransferase 1 (HPRT1) was used as the reference gene and the variations in the expression levels of target genes were determined and expressed as fold changes respect to the control, using the comparative CT method (ΔΔCT Method). The list and sequences primers used in this study can be found in Table S4.

*Western blotting:*

After treatment, the cells were washed three times with PBS and lysed in RIPA buffer (150 mM NaCl, 1% Triton X-100, 0.5% deoxycholic acid, 0.1% SDS, 20 mM Tris-HCl, pH 7.4), supplemented with protease and phosphatase cocktail inhibitor (Roche). The lysates were clarified by centrifugation, quantified using a BCA Protein Assay kit (Pierce), and resolved by SDS-PAGE and immunoblot. Blots were incubated with ECL for 3 min and exposed to x-ray films, which were then scanned. The intensity of the bands was quantitated using ImageJ.

*Immunoprecipitation and peptide pull down assay:*

For co-immunoprecipitation experiments, HeLa cells were transfected with LCS-HA, cultured for 24 hr and then lysed on ice with immunoprecipitation buffer (150 mM NaCl, 1% Igepal, 20 mM Tris-HCl, pH 7.4, protease inhibitors). The lysate was cleared and quantitated using the BCA Protein Assay kit (Pierce). Equal amounts of proteins were incubated with magnetic beads conjugated with anti-HA antibody (Sigma) overnight at 4 C. Then, the samples were washed extensively with immunoprecipitation buffer, eluted with HA peptide (Sigma), and analyzed by SDS-PAGE and immunoblotting with anti-HA and anti-GOLPH3 antibodies (see Table S5).

Biotinylated N-terminal amino acid tails of Golgi enzymes were resuspended in 150 mM NaCl and 20 mM Tris pH 7.4. For each sample, between 1 and 5 µg of peptide was incubated with 40 µL of pre-washed monomeric avidin beads (Pierce) and incubated 1 hr at 4 C. Beads functionalized with CTs were mixed with 500 µg of HeLa cell lysate (lysis buffer contains 150 mM NaCl, 20 mM Tris HCl pH 7.4, 0.1 % triton X-100, 1 mM DTT, protease and phosphatase inhibitors, and 5 mM EDTA) or 5 µg of purified ΔGOLPH3-His and incubated overnight at 4 C. Then, the samples were washed extensively with lysis buffer and analyzed by SDS-PAGE and immunoblotting with anti-His or anti-GOLPH3 antibody (see Table S5).

*Immunofluorescence and confocal microscopy:*
Cells grown on coverslips were fixed with 4% paraformaldehyde and permeabilized with 0.2% saponin (as described in (Trucco et al., 2004)). Samples were then incubated with selected antibodies against the antigen of interest at 4 C overnight followed by second antibodies labeled with Alexa Fluor dyes (Invitrogen). The coverslips were then mounted and analyzed under a confocal microscope (LSM700; Carl Zeiss; 40x or 63× oil-immersion objective (1.4 NA)). Images were processed using Metamorph 7.7.3.0 (Universal Imaging) and ImageJ software. Excel and GraphPad Prism version 5.0 software were used for data analyses and graphing. Adobe Photoshop CS3 was used to adjust the contrast of the images (for presentation only), whereas Adobe Illustrator CC 2014 (Adobe Systems) was used to illustrate figures and draw models.

*Live cell imaging:*
For live cell imaging, HeLa-M cells transiently expressing LCS-GFP_RUSH were grown on glass-bottomed 35-mm dishes which were then mounted on a Zeiss LSM700 laser scanning microscope under controlled temperature and $CO_2$. The images were acquired at regular interval (488 for excitation; PMT: 510–550 nm; 512×512 pixels; frame average, 4).

*Immuno-electron microscopy:*

The samples were fixed and processed as described previously (3). In brief, HeLa cells were fixed with 4% paraformaldehyde and 0.05% gluteraldehyde for 30 min at room temperature, then washed with PBS/0.02 M glycine and embedded in 12% gelatin in PBS. Small blocks (1 mm) were obtained and infiltrated with 2.3 M sucrose at 4 C. Ultrathin cryosections (50–70 nm) were prepared using a UC7 Leica cryo-ultramicrotome and incubated with indicated antibodies of interest followed by protein A gold. Random sampling of Golgi stacks was performed using a Tecnai-12, FEI transmission electron microscope, and pictures were acquired using a Veletta CCD digital camera. Morphological (clathrin at the TGN) and compositional (cis Golgi marker GM130) criteria were used to define the polarity of the Golgi stacks. For quantitation (performed with iTEM image acquisition software) cis, medial or trans Golgi were defined as previously described (Rizzo et al., 2013). Cis indicated the cis-most cisterna in the case of a stack with three or four cisternae, and the two cis-most cisternae in the case of a stack with five cisternae (here, the LD was the mean of the two).

Trans was the last (trans) cisterna. Medial was the remaining one or two central cisternae. Vesicles were round profiles of 50–80 nm in diameter, present within 200 nm of the rims of the stack. The distribution of LCS-HA, GalT1 and GOLPH3 within the Golgi stacks was expressed as linear density (LD), which represents the number of gold particles/µm.

*GOLPH3 cloning into pT-7.7 and pCDNA4B:*

To clone GOLPH3 into the pT-7.7 expression vector, residues 1–57 were substituted with a 6xHis-tag and the fragment was amplified using vector/GOLPH3 as a template, recombinant Taq DNA polymerase High Fidelity (Vent DNA polymerase; NEB, UK), and oligonucleodides 5′-GOLPH3His (5′-actttaagaaggagatatacatatgcatcatcatcatcatcataccaggctgaccctga-3′) and 3′-GOLPH3Stop (5′-caggtcgactctagaggatccttacttggtgaacgccgc-3′), as forward and reverse primer (over-lapping ends with the destination vector indicated in bold), in a 35-cycle PCR (95 C; 30 sec, 48 C; 30 sec, 72 C; 1 min). Primers were designed to clone GOLPH3 in-frame with the expression elements of the vector. z, the PCR product was used as a mega-primer in a second PCR for gene cloning. The RF reactions were as follows: a single denaturation step (95 C, 5 min) was performed followed by 20 cycles PCR (95 C, 1 min; 50 C, 30 s; 72 C, 5 min), and a final elongation step of 10 min at 72 C. RF reactions were performed in a final volume of 50 µL including the following components: 50 ng of target DNA, 500 ng of PCR product (mega-primers), 200 µM of each dNTP, 1x Pfu buffer, and 5U Pfu Turbo Cxhotstart DNA polymerase (Agilent). A 30 µL aliquot of each PCR mix was digested using 20 U DpnI (NEB, UK) (37 C, 1 hr), to digest the parental plasmid DNA, and 20 µL of each aliquot was used to transform Escherichia coli Top10 strain (Invitrogen). The same procedure was used to clone GOLPH3-[illegible]57 into pCDNA4B, with forward and reverse oligonucleotides 5′-GOLPH3pCDNA4B (5′-GCTTGGTACCGAGCTCGGATCCATGACCAGGCTGACCCTGATGGAGGAAG-3′) and 3′-GOLPH3pCDNA4B (5′-ACTGTGCTGGATATCTGCAGAATTCTTACTTGGTGAACGCCGCCACCA-3′), respectively (ends overlapping with the destination vector in bold). Colony PCR screening was used to search positive clones. The cloned fragments were completely sequenced to verify that during the amplification procedure undesired mutations had not been introduced.

*GOLPH3 over-expression and purification:*

Protein expression was performed in 2 L LB medium containing 100 µg/mL ampicillin at 37 C; when the cell culture reached the optical density (OD) of 0.6 (at 600 nm) protein expression was induced by IPTG (0.5 mM). After 2 hr, cells were harvested by centrifugation (3,000 x g, 4 C, 10 min). All subsequent procedures were carried out at room temperature, unless otherwise indicated. Wet frozen cells (10 g) were thawed and dissolved in 50 mL buffer containing 20 mM phosphate pH 7.5; 50 mM NaCl; and protease inhibitor cocktail (Sigma-Aldrich). Cell disruption was obtained by ultrasound apparatus (Branson), using 5 cycles at 40 % power for 30 sec. Cell debris was removed by centrifuging (30,000 x g, 4 C, 20 min). Then, 2 mL of Ni-NTA resin (Qiagen) was added to the soluble fraction and incubated for 16 hr at 4 C under gentle stirring. To reduce non-specifically bound proteins to the resin, imidazole was added at final concentration of 10 mM. The resin was loaded onto a column (20 mL), then washed with 20 mL phosphate buffer containing NaCl and imidazole (10 mM); two additional wash steps were performed by using phosphate buffer containing imidazole at 20 and 50 mM and eluted with phosphate buffer containing 200 mM imidazole; the sample was recovered in fractions of 1 mL. Fractions were pooled (about 10 mL total volume), concentrated by ultrafiltration onto a 10,000 cut-off cellulose membrane (Amicon) to 4 mL, divided in 2 mL aliquots, and loaded onto a Hi-Load 16/60 Superdex 75 column (GE Healthcare). The column was equilibrated and eluted with 20 mM phosphate buffer containing 50 mM NaCl and 5 mM imidazole. The flow rate was 0.5 mL/min. The presence of GOLPH3 was monitored by SDS-PAGE; fractions containing the protein were pooled and concentrated by ultrafiltration up to 2 mL; 2 mg pure enzyme was purified from 10 g of frozen cells.

*Electrophoretic analysis:*
SDS-PAGE analysis (12.5%) was performed as described by (Laemmli, 1970), at room temperature. The "10 kDa-250 kDa Precision plus protein standard" (Bio Rad) was used as a molecular weight standard.

*Proliferation assays:*
For in adherence colony formation assays, after treatment, 250 cells for each condition were seeded into 6-well plates and grown for 10 days. Cell colonies were glutaraldehyde-fixed and stained with crystal violet (0.1 % w/v) for 10 min, washed with water and counted using ImageJ software. Soft-agar growth assays were performed on 12-well plates in triplicate. For each well, 1 x 104 cells were mixed thoroughly in cell growth medium containing 0.3% agarose (Invitrogen, 16500-500) in DMEM plus 10% FCS, 1% penicillin/streptomycin, and 0.5 mM sodium pyruvate, followed by plating onto bottom agarose prepared with 0.6% agarose in DMEM and 10% FCS. The wells were allowed to solidify and 200 µL of growth medium was added on top and refreshed every 4 or 5 days to avoid agar drying. After 10–30 days, depending on the cell line, colonies were stained with nitrotetrazolium blue chloride (Sigma N6876), scanned, and counted using ImageJ.

*Flow cytometry analysis:*
After treatment, HeLa cells were subjected to trypsin digestion, fixed with 4% paraformaldehyde, washed, and resuspended with PBS. To visualize cell-surface-specific carbohydrates, cells were incubated with bacterial toxins or lectins for 1 hr at 4 C. Then, cells were extensively washed with PBS and incubated with fluorescence-labeled secondary antibodies when required, or directly analyzed by BD FACS ARIAIII cell sorter (BD Biosciences). Cells incubated with secondary antibody alone, or unlabeled cells, were used as a negative control. The cell-surface expression of sugars and GSLs of selected cells were further analyzed in the gated region. Antibodies, lectins and toxins used are described in (Table S5). To visualize the PM localization of luminal GFP chimeras containing the transmembrane domain of the sucrase–isomaltase (see Fig. 1D, E) and the relative N-terminal

LCS cytosolic tails, HeLa cells transiently expressing the constructs were subjected to trypsin digestion, washed and resuspended with PBS/0.5 % BSA. Cells were incubated with anti-GFP antibodies, respectively (without permeabilization) for 1 hr at 4 C. Then, cells were extensively washed with PBS/0.5 % BSA and incubated with fluorescence-labeled secondary antibodies Alexa Fluor 568 and analyzed by BD FACS ARIAIII cell sorter (BD Biosciences).

*Yeast strains, plasmids and culture conditions:*

Yeast strains and primers used in this study are listed in the tables. Yeast Strains expressing Mnn9-mCherry, Mnn9-sfGFP, Vph1-sfGFP, Vph1-mCherry were constructed by a PCR-based method using pFA6a plasmids as a template (Kurokawa et al., 2019) (primers 5'-atttggcttaccaaactatttggtttatcacatagaggaagagaaccatcggatccccgggttaattaa-3' and 5'-attatctttcaataacgctatagcttctgtatgctttttgctcagttgcgaattcgagctcgtttaaac-3' for Mnn9-mCherry and Mnn9-sfGFP and primers 5'-ataaagacatggaagtcgctgttgctagtgcaagctcttccgcttcaagccggatccccgggttaattaa-3' and 5'-tatttaatgaagtacttaaatgtttcgcttttttaaaagtcctcaaaatgaattcgagctcgtttaaac-3' for Vph1-mCherry and Vph1-sfGFP). Sys1-GFP and Sec7-GFP were expressed under the control of the *ADH1* promoter on the low-copy plasmid pRS316 (Kurokawa et al., 2019).

*Live cell imaging of yeast:*

Yeast cells were grown in MCD medium [0.67% yeast nitrogen base without amino acids, 0.5% casamino acids (Difco Laboratories Inc.), and 2% glucose] with appropriate supplements. For live imaging, cells were grown to a mid-log phase at 24 C. Cells were immobilized on glass slides using concanavalin A and imaged by super-resolution confocal microscopy (SCLIM). SCLIM was developed by combining Olympus model IX-71 inverted fluorescence microscope with a UPlanSApo 100 X NA 1.4 oil objective lens (Olympus, Japan), a high-speed and high-signal-to noise-ratio spinning-disk confocal scanner (Yokogawa Electric, Japan; (Kurokawa et al., 2014), a custom-made spectroscopic unit, image intensifiers (Hamamatsu Photonics, Japan) equipped with a custom-made cooling system, magnification lens system for giving 266.7 X final magnification, three CW lasers (473 nm, 561 nm and 671 nm), and three EM-CCD cameras (Hamamatsu Photonics, Japan) for green, red, and infrared observation. Image acquisition was executed by a custom-made software (Yokogawa Electric, Japan). For 3D and 4D live imaging, we collected optical sections spaced 100 nm or 200 nm apart in stacks by oscillating the objective lens vertically with a custom-made piezo actuator. Z stack images were converted to 3D voxel data and processed by deconvolution with Volocity (Perkin Elmer, MA) using the theoretical point-spread function for spinning-disk confocal microscopy. For the photobleaching analysis, sfGFP tagged proteins in whole area was photobleached with 473 nm CW laser at 90% of the laser power. Before and after photobleaching, we took 3D images of sfGFP tagged proteins with 473 nm CW laser 10 % of laser power at indicated time. MetaMorph software (Molecular Devices, CA) was used for presenting time-lapse images and analyzing fluorescence signal changes for cisternal conversion and quantifying fluorescence intensities in selected cells before and after photobleaching. For FRAP analysis, we conducted 2D time-lapse observation at every 2 minutes before and after photobleaching and analyzed fluorescence intensities in selected cells after background subtraction by MetaMorph software.

*Experimental conditions for LCS-GFP_RUSH traffic assay:*
i) GOLPH3-KD HeLa cells were transiently transfected with LCS-GFP_RUSH at 37 C in absence of biotin. Cells were then treated with nocodazole (33 µM) and cycloheximide (50 mg/mL) for 3 hours prior to shifting the temperature to 10 C for 60 min in the presence of biotin (40 µM). Then, cells were re-shifted to 37 C for the indicated times (Table S6), fixed and subjected to IF. See Table S6 for a scheme of the synchronization protocol.
ii) GOLPH3-KD HeLa cells were transiently transfected with LCS-GFP_RUSH and culture at 37 C in the absence of biotin for 24 hours. Cells were then infected with VSV-G and cultured at 40 C for 3 hours in presence of nocodazole (33 µM), prior to shifting the temperature to 10 C for 60 min in presence of biotin (40 µM) and cycloheximide (50 mg/mL). Then, cells were re-shifted to 32 C for the indicated time, fixed and subjected to IF. See Table S6 for a scheme of the synchronization protocol.

Lipid Analysis

*(Glyco)sphingolipid measurements:*

*HPLC-Mass Spectrometry:*
Glycophingolipids were analyzed by liquid chromatography tandem mass spectrometry (LC-MS/MS) as described earlier (Bielawski et al., 2006). Briefly, extracts were analyzed with a Quantum Ultra triple quadrupole mass spectrometer connected to an Accela HPLC and Accela autosampler using a solvent gradient. Ceramides identity was achieved through MRM analysis with soft fragmentation. Quantitative analysis is based on calibration curves generated for each lipid. Levels of SLs were normalized to inorganic phosphate (Pi) released from total phospholipids.

*HPTLC:*
Cells were pulse-labelled with 0,1 µCi/mL (≈ 5nM) $^{3}$H-D-erythro-Sphingosine for 2 hours and chased for the indicated times. Subsequently, cells were harvested and processed for lipid extractions. Lipids were spotted on silica-gel high performance-TLC (HPTLC) plates (Merck, Germany), and resolved with a mixture of chloroform: methanol: water (65:25:4 v/v/v). To visualize the unlabelled standards (i.e., Cer, GlcCer, LacCer, Gb3, GM3 and SM) the TLC plates were placed in a sealed tank saturated with iodine vapours, while the radiolabelled lipids were analysed using a RITA® TLC Analyser (Raytest, Germany), and quantified using GINA® (Raytest, Germany) software analysis. The percentage of total C.P.M. associated with each sphingolipid are reported as percentage of total sphingolipids.

*MALDI MS:*

Lipid extraction

Total lipid extracts were prepared using a standard MTBE protocol followed by a methylamine treatment for sphingo- and glycosphingolipids analysis by mass spectrometry. Briefly, cell pellet was resuspended in 100 µL $H_2O$. 360 µL methanol and 1.2 mL of MTBE were added and samples were placed for 10 min on a vortex at 4 C followed by incubation for 1 h at room temperature on a shaker. Phase separation was induced by addition of 200 µL of $H_2O$. After 10 min at room temperature, samples were centrifuged at 1000 g for 10 min. The upper (organic) phase was transferred into a glass tube and the lower phase was re-extracted with 400 µL artificial upper phase [MTBE/methanol/$H_2O$ (10:3:1.5, v/v/v)]. The combined organic phases were dried in a vacuum concentrator. Lipids where then resuspended in 500 µL of $CHCl_3$ and divided in two aliquots for a further methylamine treatment. 500 µL of freshly prepared monomethylamine reagent [methylamine/$H_2O$/n-butanol/methanol (5:3:1:4, (v/v/v/v)] was added to the dried lipid extract and then incubated at 53 C for 1 h in a water bath. Lipids were cooled to room temperature and then dried. The dried lipid extract was then extracted by n-butanol extraction using 300 µL water-saturated n-butanol and 150 µL $H_2O$. The organic phase was collected, and the aqueous phase was re-extracted twice with 300 µL water-saturated n- butanol. The organic phases were pooled and dried in a vacuum concentrator. Lipids were then resuspended in 500 µL of $CHCl_3$ and analyzed by MALDI-MS. 30 mg/mL 2,5-DHB was freshly prepared in acetonitrile/water solution (50:50 v/v) with 0.1% TFA. An equivalent volume of sample solution (50 µL) was then mixed with matrix before deposition on the MALDI target. All mass spectrometry analysis for the identification of lipids (m/z 600-1800) were obtained using an Applied Biosystems 4800 MALDI-TOF/TOF mass spectrometer equipped with a 200 Hz tripled-frequency Nd:YAG pulsed laser with 355 nm wavelength. Measurements were performed in positive ion reflection mode at an accelerating potential of 20 kV. Each mass spectra were obtained by applying a laser energy of 4600 watts/cm2, averaging 4000 single laser shots/spectrum.

*Untargeted Lipidomics:*

For phospholipid analysis, lipid extracts (2 µL injection volume in $CHCl_3$:MeOH 2:1) were separated over an 8 minute gradient at a flow rate of 200 µL/min on a HILIC Kinetex Column (2.6lm, 2.1 × 50 mm2) on a Shimadzu Prominence UFPLC xr system (Tokyo, Japan). Mobile phase A was acetonitrile:methanol 10:1 (v/v) containing 10 mM ammonium formate and 0.5% formic acid while mobile phase B was deionized water containing 10 mM ammonium formate and 0.5% formic acid. The elution of the gradient began with 5% B at a 200 µL/min flow and increased linearly to 50% B over 7 min, then the elution continued at 50% B for 1.5 min and finally the column was re-equilibrated for 2.5 min. MS data were acquired in full-scan mode at high resolution on a hybrid Orbitrap Elite (Thermo Fisher Scientific, Bremen, Germany). The system was operated at 240,000 resolution (m/z 400) with an AGC set at 1.0E6 and one microscan set at 10-ms maximum injection time. The heated electrospray source HESI II was operated in positive mode at a temperature of 90 C and a source voltage at 4.0KV. Sheath gas and auxiliary gas were set at 20 and 5 arbitrary units, respectively, while the transfer capillary temperature was set to 275 C.

*Isothermal titration calorimetry (ITC):*

ITC experiments were performed in a buffer containing 300 mM NaCl, 10 mM Bicine pH 8.5 and 1 mM DTT. LCS Peptides were synthesized and delivered as lyophilized powder with a biotin moiety located at the N terminus (Charite Universitaetsmedizin Berlin, Germany). The peptides were dissolved in buffer, centrifuged at 14000 x g for 10 minutes and only the supernatant was used. The dissolved peptide concentrations were calculated based upon their absorbance at 280 nm and their corresponding molar extinction coefficient. Experiments consisted of titrations of 20 injections of 2 µL of titrant (peptides) into the cell containing GOLPH3 protein at a 25-fold lower concentration. Typical concentrations for the titrant were around 2.5 mM for experiments depending on the affinity. Experiments were performed at 25 C and a stirring speed of 1000 rpm on an MicroCal PEAQ-ITC (Malvern Panalytic). All data were processed using MicroCal PEAQ-ITC Analysis Software and fit to a one-site binding model after background buffer subtraction.

*TMA Building:*

72 surgical specimens from Non Small Cell Lung Cancer (NSCLC) patients collected from 2006 to 2010, at the National Cancer Institute "Giovanni Pascale" of Naples, were used for building a tissue microarray (TMA). The TMA was built using two cores from different areas selected on the H&E-stained slides and, whenever possible, one core of normal tissue of the same tissue block. Tissue cylinders with a diameter of 1 mm were punched from morphologically representative tissue areas of each "donor" tissue block and brought into one recipient paraffin block (3 × 2.5 cm) using a semi-automated tissue arrayer (Galileo TMA).

*Immunohistochemistry:*

A standard protocol was used for the immunostaining of the paraffin embedded samples to evaluate the expression of GOLPH3 and LCS; an appropriate external positive control tissue was used for each staining procedure; the negative control consisted of performing the entire IHC procedure on an adjacent section in the absence of the primary antibody. Briefly, paraffin slides (4 µm) were cut, deparaffinized in xylene and rehydrated through graded alcohols. Antigen retrieval was performed according to manufacturer's instructions, with slides heated in high pH solution (Dako EnVision FLEX Target Retrieval Solution 50x) for LCS and in low pH solution (Dako EnVision FLEX Target Retrieval Solution 50x) for GOLPH3, in a bath for 20 min at 97 C. The endogenous peroxidase was inactivated with 3% hydrogen peroxide and then the protein block (BSA 5% in PBS 1×) was performed. TMAs sections were incubated with the primary antibodies according to the specific conditions tested: anti-GOLPH3 (Polyclonal Rabbit-, Abcam Cat # ab98023 RRID: AB_10860828 - dilution 1:50, incubated 1 h) and anti-LCS (B4GALT5) (Monoclonal Mouse antibody- Clone 6D4- Kind gift from Henrik Clausen, Denmark - incubated 1 h). The tissue sections were incubated with IgG biotinylated secondary antibodies for 40 min at room temperature. Immunoreactivity was visualized by means of avidin-biotin-peroxidase complex kit reagents (Novocastra, Newcastle, UK) as the chromogenic substrate. Finally, the sections were developed with diaminobenzidine and counterstained with hematoxylin. Stained slides were analysed using an optical microscope (Olympus BX41). Expressions of the biomarkers were evaluated semiquantitatively based on the staining intensity and the number of immunoreactive cells. First, cytoplasmic-staining intensity was scored as follows: no staining (score 0), weak expression cytoplasmatic staining (score 1), moderate expression cytoplasmatic staining (score 2), high expression cytoplasmatic staining (score 3). Second, the percentage of positive cells was scored: no positive cells (0), 10% positive cells or less (1), 11% to 50% positive cells (2), 51% to 75% positive cells (3), and more than 75% positive cells (4). The immunoreactivity scores of the cancer tissue samples were determined based on the staining intensity and area of positive staining according to the method used by Wang et al. (Wang et al., 2015).

*Fish Assay:*

FISH for GOLPH3 status was performed on TMA section using a mix of two probes according to the manufacturer's instructions, one labelled Orange to detect GOLPH3 (5p13.3) and ones Green to Chromosome 5 control probe (5p11) (Empiregenomics). Briefly, TMA was cut in 5 µm thick sections. The slides were baked for 1 hour at 60 C and then placed in a solution for deparaffination followed by dehydration steps. The slides were treated with protease solution and pretreatment with a commercial kit (Paraffin pretreatment reagent kit; Vysis) on the half-automated VP2000 processor system (Abbott Molecular, Wiesbaden, Germany). After pretreatment, the slides were denatured with the mixture of probes for 5 min at 85 C and hybridized at 37 C overnight in a Thermobrite (Leica). Post-hybridization 2XSSC 0.1%NP40 washes were performed both at room temperature and 72 C, then, the slides were stained with DAPI before analysis. Normal tissue and lymphocytes were taken into account as internal controls. Tumor core tissue was entirely scanned for amplification by using a 63 objective and appropriate filter sets (BX61 fluorescent microscope; Olympus). If GOLPH3 signal showed a homogenous distribution, random areas were used for counting the signals. Each FISH slide was evaluated by 2 independent investigators in a blind manner. A minimum of 40 nuclei were counted on each spot. Amplification of GOLPH3 was defined when GOLPH3/Centromere 5 ratio was ≥ 2.0 or when the average of GOLPH3 signals for cell was ≥6.0. The copy number gain was considered when the average of GOLPH3/cell signals were >4.0 and <6.0.

*Quantification and Statistical Analysis:*

Error bars correspond to either standard deviation (S.D.) or standard error of the mean (S.E.M.) according to the different experiments and as indicated in the figure legends. Statistical evaluations report on Student's t test *$p < 0.05$, **$p < 0.01$, and ***$p < 0.001$ (ns, not significant).

Supplementary Tables.

**Table S1** TMA FISH and IHC analysis. N/A, data not available; CGN, copy number gain (i.e., cases where the average of GOLPH3/cell FISH signals were >4.0 and <6.0); AMPL, amplified i.e., cases where GOLPH3/Centromere 5 ratio was ≥ 2.0 or when the average GOLPH3 FISH signals per cell were ≥6.0).

| sex | age | FISHGOLPH3 | GOLPH3:IHC | LCS:IHC |
|---|---|---|---|---|
| M | 70 | NO | high | weak |
| F | 62 | CNG | high | med |
| M | 75 | CNG | med | med |
| F | 76 | N/A | med | med |
| M | 67 | CNG | weak | weak |
| M | 72 | CNG | med | neg |
| M | 44 | NO | neg | neg |
| F | 65 | NO | weak | weak |
| M | 65 | NO | med | med |
| M | 64 | NO | med | neg |
| F | 51 | AMPL | neg | neg |
| M | 48 | NO | med | med |
| M | 63 | CNG | high | weak |
| M | 57 | N/A | neg | neg |
| F | 57 | NO | med | weak |
| M | 78 | AMPL | neg | neg |
| M | 75 | NO | med | med |
| M | 54 | NO | med | med |
| M | 47 | NO | med | neg |
| M | 56 | CNG | high | weak |
| F | 54 | NO | high | weak |
| M | 58 | NO | med | med |
| F | 66 | N/A | neg | neg |
| M | 63 | NO | high | weak |
| F | 66 | AMPL | med | med |
| F | 68 | NO | med | weak |
| M | 71 | NO | neg | neg |
| M | 59 | NO | med | med |
| M | 56 | CNG | med | med |
| M | 63 | NO | med | med |
| M | 52 | CNG | med | med |

| | | | | |
|---|---|---|---|---|
| M | 63 | AMPL | weak | weak |
| M | 78 | NO | med | med |
| M | 69 | CNG | med | med |
| M | 78 | NO | high | med |
| M | 71 | AMPL | med | med |
| M | 65 | NO | med | med |
| M | 59 | NO | weak | weak |
| M | 56 | NO | neg | neg |
| M | 59 | NO | med | med |
| M | 55 | NO | med | med |
| M | 67 | NO | high | med |
| M | 70 | NO | weak | med |
| M | 72 | AMPL | weak | weak |
| M | 55 | NO | med | med |
| M | 60 | NO | med | neg |
| F | 47 | AMPL | med | neg |
| F | 62 | NO | weak | weak |
| F | 55 | NO | weak | weak |
| M | 64 | NO | weak | weak |
| M | 68 | NO | weak | med |
| M | 54 | CNG | med | med |
| M | 77 | NO | med | weak |
| F | 71 | CNG | high | high |
| M | 63 | NO | weak | weak |
| F | 54 | NO | weak | med |
| F | 53 | CNG | high | med |
| F | 59 | CNG | med | med |
| M | 57 | CNG | high | med |
| F | 69 | NO | med | med |
| F | 72 | CNG | med | weak |
| F | 70 | NO | med | weak |
| F | 66 | NO | high | neg |
| M | 75 | NO | high | med |
| M | 53 | NO | med | neg |
| M | 67 | CNG | neg | med |
| F | 52 | CNG | neg | med |
| M | 73 | NO | weak | neg |
| M | 65 | AMPL | med | med |
| M | 71 | NO | weak | weak |
| M | 68 | CNG | neg | neg |

| | | | | |
|---|---|---|---|---|
| F | 28 | NO | high | high |

**Table S2** List of siRNAs used in this study

| Human Gene | Accession Number | siRNA Sequence |
|---|---|---|
| GOLPH3 | NM_022130 | 1# 5'-aaaugauguguaacccucgcggucc-3'<br>2 #5'-aauccagaugauauacagucauucc -3'<br>6 #5'-ggagaggaagguuacaacua-3'<br>7# 5'-ucaaggaccgcgaggguua -3' |
| B4GALT5 (LCS) | NM_004776 | 1# 5'-gugaaauuggaauggauua-3'<br>2# 5'-gcuuaacaguggaacaauu-3'<br>4# 5'-ggaaagugaucgcaacuau-3'<br>5# 5'-gaaagacucccuuccauga-3' |
| SMS1 | NM_147156 (from Capasso S. et al.) | 1# 5'-cuacacucccaguaccugg-3'<br>2 #5'-cacacuauggccaaucagcaa-3' |
| Gb3S | NM_017436 (from Russo D. et al.) | 1# 5'-agaaagggcagcucuauaauu-3'<br>2# 5'-ggacacggacuucauuguuuu-3'<br>3# 5'-ugaaagggcuuccgggugguu-3'<br>4# 5'-gcacucauguggaaguucguu-3' |

**Table S3** List of biotinylated peptides used in this study

| Name | Peptide sequence |
|---|---|
| LCS-tail-WT | MRARRGLLRLPRRSLLA-Biot |
| LCS-tail-mut1 | MRARRGLLAAAAASLLA-Biot |
| LCS-tail-mut2 | AAAAAGLLRLPRRSLLA-Biot |
| LCS-tail-mut3 | MRARRAAAAAPRRSLLA-Biot |
| GalT1-tail-wt | MRLREPLLSGSAAMPGA-Biot |
| Gb3S-tail | MSKPPDLLLRLLRGAPRQRVCT-Biot |
| GM3S-tail | MRRPSLLLKD-Biot |
| B4GALNT1-tail | MWLGRRA-Biot |
| GD3S-tail | MSPCGRARRQTSRGAMAVLAWKFPRTRLP-Biot |
| LC3S-tail | MRMLVSGRRVKKWQLIIQLF-Biot |
| FUT5-tail | MDPLGPAKPQWLWRR-Biot |
| B3GNT4-tail | MLPPQPSAAHQGRGGRSGLLPKGPAMLC-Biot |
| GALNT3-tail | MAHLKRLVKLHIKRHYHKK-Biot |

**Table S4** List of primers used in this study

| Human | Forward Primer | Reverse Primer |
|---|---|---|
| GOLPH3 | 5’-cccaagcttatgacctcgctgacccagc-3’ | 5’- ggggtacccttggtgaacgccgcc-3’ |
| LCS | 5’-caatcggtgctcaggtttatg-3’ | 5’-ggtttcactgtggttcaagtc-3’ |
| HPRT1 | 5’-agcttgctggtgaaaaggac-3’ | 5’-gtcaagggcatatccaacaac-3’ |
| Yeast | Forward Primer | Reverse Primer |
| MNN9 | 5’-atttggcttaccaaactatttggtttatcacatagaggaagagaaccatcggatcccgggttaattaa-3’ | 5’-attatctttcaataacgctatagcttctgtatgctttttgctcagttgcgaattcgagctcgtttaaac-3’ |
| VPH1 | 5’-ataaagacatggaagtcgctgttgctagtgcaagctcttccgcttcaagccggatccccgggttaattaa-3’ | 5’-tatttaatgaagtacttaaatgtttcgcttttttaaaagtcctcaaaatgaattcgagctcgtttaaac-3’ |

**Table S5** List of antibodies, lectins and toxins used in this study for immunofluorescence (IF), Western Blotting (WB), cryo-Electron Microscopy (cryo-EM), and FACS.

| REAGENT or RESOURCE | SOURCE | IDENTIFIER |
|---|---|---|
| Polyclonal Rabbit GOLPH3 | Abcam | Cat # ab98023<br>RRID: AB_10860828 |
| Polyclonal Mouse GOLPH3 | Abcam | Cat # ab69171<br>RRID:AB_2279272 |
| Monoclonal Mouse GAPDH | Santa Cruz Biotechnology | Clone 6C5<br>Cat #sc-32233<br>RRID:AB_627679 |
| Monoclonal Mouse HA-Tag | BioLegend/Covance | Clone 16B12<br>Cat #MMS-101P<br>RRID:AB_10063630 |
| Monoclonal Rabbit HA-Tag | Cell Signaling Technology | Clone C29F4<br>Cat #3724 |
| Polyclonal Rabbit LAMPI | Abcam | Cat # ab24170<br>RRID: AB_775978 |
| Polyclonal Rabbit B4GALT1 | Sigma | Cat # HPA010807<br>RRID: AB_1078254 |
| Polyclonal Sheep anti- | BioRad/AbD-Serotec | Cat #AHP500G |

| | | |
|---|---|---|
| human TGN46 | | RRID:AB_323104 |
| Monoclonal Mouse GM130 | BD Biosciences | Clone 35<br>Cat # 610822<br>RRID: AB_398141 |
| Monoclonal Mouse GFP | Abcam | Cat #ab6556<br>RRID:AB_305564 |
| Monoclonal Mouse GFP | Abcam | Cat #ab1218<br>RRID:AB_298911 |
| Monoclonal Mouse anti-β Actin | Sigma | Clone AC-74<br>Cat #A2228<br>RRID:AB_476697 |
| Monoclonal Mouse B4GALT5 (LCS) | Kind gift from Henrik Clausen, Denmark | Clone 6D4 |
| Monoclonal Mouse B4GALT5 (LCS) | Kind gift from Henrik Clausen, Denmark | Clone 1E10 |
| Polyclonal Rabbit B4GALT5 (LCS) | ThermoFisher Scientific | Cat PA5-25282<br>RRID:AB_2542782 |
| Monoclonal Mouse Pentahistidine | ThermoFisher Scientific | Cat P21315<br>RRID:AB_1500376 |
| Polyclonal Rabbit Phospho-Akt (Ser473) | Cell Signaling Technology | Cat #9271<br>RRID:AB_329825 |
| Polyclonal Rabbit Phospho-Akt (Thr 308) | Abcam | Cat# ab66134<br>RRID:AB_1141017 |
| Polyclonal Rabbit Akt | Cell Signaling Technology | RRID:AB_329827 |
| p21 Waf1/Cip1 Rabbit monoclonal antibody | Cell Signaling Technology | Clone 12D1<br>Cat #2947<br>RRID:AB_823586 |
| Polyclonal Rabbit p70S6K | Cell Signaling Technology | Clone 49D7<br>Cat#2708<br>RRID:AB_390722 |
| Polyclonal Rabbit Phospho-p70S6K (Thr 389) | Cell Signaling Technology | Clone 108D2<br>Cat#9234<br>RRID:AB_2269803 |
| Polyclonal Rabbit 4E-BP1 (53H11) | Cell Signaling Technology | Cat#9644<br>RRID:AB_2097841 |
| Polyclonal Rabbit Phospho-4E-BP1 (Ser65) | Cell Signaling Technology | Clone 174A9<br>Cat#9456<br>RRID:AB_823413 |
| Monoclonal Mouse VSV-G | Sigma | SAB4200695 |

| | | |
|---|---|---|
| ShTxB-CY3 | Dr. Ludger Johannes | N/A |
| ChTxB-AlexaFluor 488 | Invitrogen | Cat # C-22841 |
| Anti-mouse, donkey Alexa Fluor 488 | ThermoFisher Scientific | Cat #A-21202 RRID:AB_141607 |
| Anti-mouse, donkey Alexa Fluor 568 | ThermoFisher Scientific | Cat #A10037 RRID:AB_2534013 |
| Anti-mouse, goat Alexa Fluor 633 | ThermoFisher Scientific | Cat #A-21052 RRID:AB_141459 |
| Anti-rabbit, donkey Alexa Fluor 488 | ThermoFisher Scientific | Cat #A-21206 RRID:AB_141708 |
| Anti-rabbit, donkey Alexa Fluor 568 | ThermoFisher Scientific | Cat #A10042 RRID:AB_2534017 |
| Anti-rabbit, goat Alexa Fluor 633 | ThermoFisher Scientific | Cat #A-21070 RRID:AB_2535731 |
| Anti-sheep, donkey Alexa Fluor 633 | ThermoFisher Scientific | Cat #A-21100 RRID:AB_2535754 |
| Anti-goat, donkey Alexa Fluor 568 | ThermoFisher Scientific | Cat #A-11057 RRID:AB_142581 |
| Anti-mouse, donkey DyLight 405 | Jackson Laboratory | Cat # 712-475-153 RRID:AB_2340681 |
| DyLight 549 Streptavidin | Jackson Laboratory | Cat # 016-500-084 |
| Biotinylated Aleuria aurantia Lectin (AAL) | Vector Laboratories | Cat #B-1395 |
| Biotinylated Sambucus nigra Lectin (SNA) | Vector Laboratories | Cat #B-1305 |
| Biotinylated Peanut Agglutinin (PNA) | Vector Laboratories | Cat #B-1075 |
| Rhodamine labeled Ricinus communis Agglutinin I (RCA I) | Vector Laboratories | Cat #RL-1082 |
| Rhodamine labeled Phaseolus vulgaris Leucoagglutinin (PHA-L) | Vector Laboratories | Cat # RL-1112 |

Table S6. Schematic representation of LCS-GFP_RUSH traffic assays

i)

**Synchronization protocol**

| | | | | | | | | | |
|---|---|---|---|---|---|---|---|---|---|
| Temperature (°C) | 37 | → | 10 | → | 37 | → | 37 | → | 37 |
| Biotin (40 μM) | - | → | + | → | + | → | + | → | + |
| Cycloheximide (50 μg/mL) | + | → | + | → | + | → | + | → | + |
| Nocodazole (33 μM) | + | → | + | → | + | → | + | → | + |
| Time | 3 hours | → | 60 min | → | 10 min | → | 20 min | → | 30 min |

ii)

Synchronization protocol

| | | | | | | | | | |
|---|---|---|---|---|---|---|---|---|---|
| Temperature (°C) | 40 | → | 10 | → | 37 | → | 37 | → | 37 |
| Biotin (40 μM) | - | → | + | → | + | → | + | → | + |
| Cycloheximide (50 μg/mL) | - | → | + | → | + | → | + | → | + |
| Nocodazole (33 μM) | + | → | + | → | + | → | + | → | + |
| Time | 3h | → | 60 min | → | 10 min | → | 20 min | → | 30 min |

Table S7 List of bacterial and viral strains used in this study

| REAGENT or RESOURCE | SOURCE | IDENTIFIER |
|---|---|---|
| Adenoviral - Type 5 (dE1/E3) of eGFP (Ad-GFP) | Vector Biolabs | Cat # 1060 |
| Adenoviral - Type 5 (dE1/E3) of human GOLPH3 (Ad-h-GOLPH3) | Vector Biolabs | Cat # ADV-210166 |
| Adenoviral - Type 5 (dE1/E3) of human B4GALT5 (Ad-h-B4GALT5-HA) | Vector Biolabs | Cat # ADV-201912 |
| Temperature sensitive strain of vesicular stomatitis virus (ts045-VSV) | Kind gift from Bruno Goud, Institut Curie, France | N/A |
| E. coli (DH5α) | Thermo Fisher Scientific | Cat #18265017 |
| E. coli (BL-21-DE3) | Thermo Fisher Scientific | Cat #C600003 |

List of chemicals used in this study

| REAGENT or RESOURCE | SOURCE | IDENTIFIER |
|---|---|---|
| Paraformaldehyde 8% | Electron Microscopy Sciences | Cat #50-259-96 |
| Glutaraldehyde 8% | Electron Microscopy Sciences | Cat #50-262-18 |
| Methanol | JT Baker | Cat #9093 |

| | | |
|---|---|---|
| Cycloheximide | Sigma | Cat #01810 |
| Protein A Sepharose CL-4B | GE Healthcare Life Sciences | Cat #17-0780-01 |
| Anti-HA magnetic beads | ThermoFisher Scientific | Cat #88836 |
| Bafilomycin A1 from Streptomyces griseus | Sigma | Cat # 88899-55-2 |
| Insulin solution human | Sigma | Cat #11061-60-0 |
| EasyTag™ EXPRESS35S Protein Labeling Mix, [35S] | Perkin Elmer | Cat #NEG772014MC |
| Sphingosine, [3-3H]-, D-erythro>97% | PerkinElmer | Cat #NET1072050UC |
| Silica-gel high performance-TLC (HPTLC) plates | Merck, Germany | Cat #1055830001 |
| Bovine Serum Albumin Fatty acid free | Sigma | Cat #A8806 |
| HA Peptide | Sigma | Cat #I2149 |
| Biotin | Pierce | Cat #29129 |
| Protein A gold 15 nm | Cell Microscopy Core, UMC Utrecht | N/A |
| Protein A gold 10 nm | Cell Microscopy Core, UMC Utrecht | N/A |
| Protein A gold 5 nm | Cell Microscopy Core, UMC Utrecht | N/A |
| TransIT-LT1 | Mirus | Cat #MIR 2305 |
| UltraPureTM Agarose | Invitrogen | Cat #16500-500 |
| Lipofectamine LTX with PLUS | ThermoFisher Scientific | Cat #15338100 |
| Oligofectamine | ThermoFisher Scientific | Cat #12252011 |
| DMEM | Gibco/ ThermoFisher Scientific | Cat #41965 |
| RPMI 1640 | Gibco/ ThermoFisher Scientific | Cat #21875 |
| FCS | Gibco/ ThermoFisher Scientific | Cat #10437036 |
| DMEM w/o cysteine and methionine | Gibco/ ThermoFisher Scientific | Cat #21013 |
| Sodium Pyruvate (100 nM) | Gibco/ ThermoFisher Scientific | Cat #11360-070 |

| | | |
|---|---|---|
| Yeast nitrogen base without amino acids | Difco Laboratories Inc | Cat # 291940 |
| Casamino acids | Difco Laboratories Inc | Cat # 228830 |

Table S8 List of commercial assays and kits used in this study

| REAGENT or RESOURCE | SOURCE | IDENTIFIER |
|---|---|---|
| RNA easy mini | Qiagen | Cat #74106 |
| QuantiTect Reverse Transcription Kit | Qiagen | Cat #205311 |
| SYBR™ Green PCR Master Mix | ThermoFisher Scientific | Cat #4309155 |

List of recombinant DNAs used in this study

| REAGENT or RESOURCE | SOURCE | IDENTIFIER |
|---|---|---|
| GOLPH3 (untagged)-Human Golgi phosphoprotein 3 | OriGene | Cat #SC112810 |
| B4GALT1-HA | Kind gift from Hans Bakker, Germany | N/A |
| B4GALNT1-HA | GenScript | N/A |
| ST8SIA1-HA (or GD3S-HA) | GenScript | N/A |
| B3GNT5-HA | GenScript | N/A |
| B4GALT4-HA | GenScript | N/A |
| SMS1-HA | Kind gift from Giovanni D'Angelo | N/A |
| GCS-HA | Kind gift from Giovanni D'Angelo | N/A |
| Gb3S-HA | Kind gift from Antonella De Matteis | N/A |
| GM3S-HA | Kind gift from Antonella De Matteis | N/A |
| LCS-HA | This study | N/A |
| Str-KDEL_B4GALT5-SBP-EGFP (referred as LCS-GFP_RUSH) | This study. Gaelle Boncompain | N/A |
| Str-KDEL_GalTfl-SBP-EGFP (referred as B4GALT1-GFP_RUSH | Kind gift from Gaelle Boncompain (*Boncompain et al Nat Methods. 2012*) | N/A |

| | | |
|---|---|---|
| Sucrose-Isomaltase (referred as SI-GFP) | Kind gift from Stuart Kornfeld | N/A |
| B4GALT5_Met-7Ala_MutSI(TM) (referred as LCS-SI-GFP) | GenScript | N/A |
| B4GALT5_RARR_Mut-MutSI(TM) (referred as LCS-SI-RxR) | GenScript | N/A |
| B4GALT5_LXX(R/K)_Mut-SI(TM) (referred as LCS-SI-LxxR | GenScript | N/A |
| B4GALT5_RARR_LXX(R/K)_Mut-SI(TM) (referred as LCS-SI-RxR-LxxR | GenScript | N/A |

**Table S9** List of software used in this study

| REAGENT or RESOURCE | SOURCE | IDENTIFIER |
|---|---|---|
| ImageJ | NIH | https://imagej.nih.gov/ij/ |
| MetaMorph | Molecular Devices | https://www.moleculardevices.com/systems/metamorph-research-imaging |
| Prism | Graphpad | https://www.graphpad.com/scientific-software/prism/ |
| Adobe Illustrator | Adobe | www.adobe.com/products/illustrator/free-trial-download.htm |
| Adobe Photoshop | Adobe | www.adobe.com/products/photoshop.html |
| Soft Imaging service (Electron microscope) | Olympus | www.olympus-sis.com/corp/2256.htm |
| iTEM | EMSIS GmbH | https://www.emsis.eu/products/software/item/ |
| Volocity | Perkin Elmer | https://www.perkinelmer.com/lab-products-and-services/cellular-imaging/volocity-3d-visualization.html |

**Table S10** List of cell lines used in this study

| REAGENT or RESOURCE | SOURCE | IDENTIFIER |
|---|---|---|
| HeLa-M | ATCC | RRID:CVCL_R965 |
| Primary Human Fibroblasts | This study; Telethon Biobank ((http://www.telethon.it/en/funding-research/funded-projects/details/telethon-network-of-genetic-biobanks_0) | N/A |
| LNCaP | ATCC | RRID:CVCL_0395 |
| DU-145 | ATCC | RRID:CVCL_0105 |
| HeLa-mCAT#8 | Kind gift from Kentaro Hanada | N/A |
| TALEN UGCG-KO | Kind gift from Kentaro Hanada | N/A |
| TALEN LCS-KO | Kind gift from Kentaro Hanada | N/A |

**Table S11** List of yeast strains used in this study (see (Sikorski and Hieter, 1989))

| Strain | Genotype | Source |
|---|---|---|
| YPH499 | *MAT a ura3-52 lys2-801 ade2-101 trp1-Δ63 his3-Δ200 leu2-Δ1* | 1 |
| YMO132 | *YPH499ADE2::pRS402 MNN9-mCherry::natNT2* | 2 |
| YMI125 | *YPH499 ADE2::pRS402 vps74::kanMX4 MNN9-mCherry::natNT2* | This study |
| YKK935 | *YPH499 ADE2::pRS402 MNN9-sfGFP::HIS3MX6* | This study |
| YKK937 | *YPH499 ADE2::pRS402 vps74::kanMX4 MNN9-sfGFP::HIS3MX6* | This study |
| YKK987 | *YPH499 ADE2::pRS402 MNN9-sfGFP::HIS3MX6 VPH1-mCherry::natNT2* | This study |
| YKK989 | *YPH499 ADE2::pRS402 vps74::kanMX4 MNN9-sfGFP::HIS3MX6 VPH1-mCherry::natNT2* | This study |
| YKK990 | *YPH499 ADE2::pRS402 VPH1-sfGFP::HIS3MX6* | This study |
| YKK992 | *YPH499 ADE2::pRS402 vps74::kanMX4 VPH1-sfGFP::HIS3MX6* | This study |
