## Supplementary material for "The Glyco-enzyme adaptor GOLPH3 Links Intra-Golgi Transport Dynamics to Glycosylation Patterns and Cell Proliferation": Sipplementary Figures

### Supplementary Figure 1

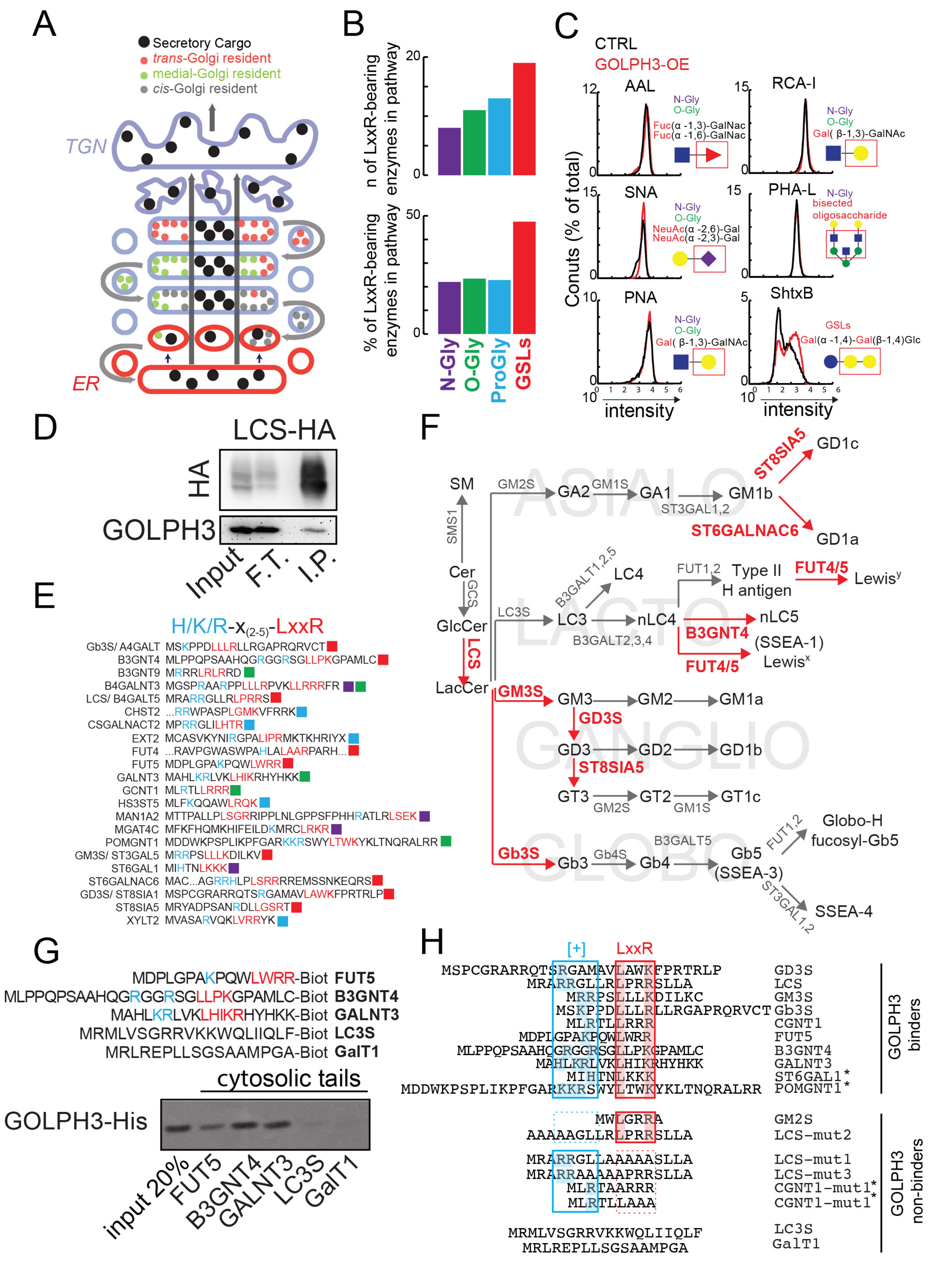

### Supplementary Figure 2

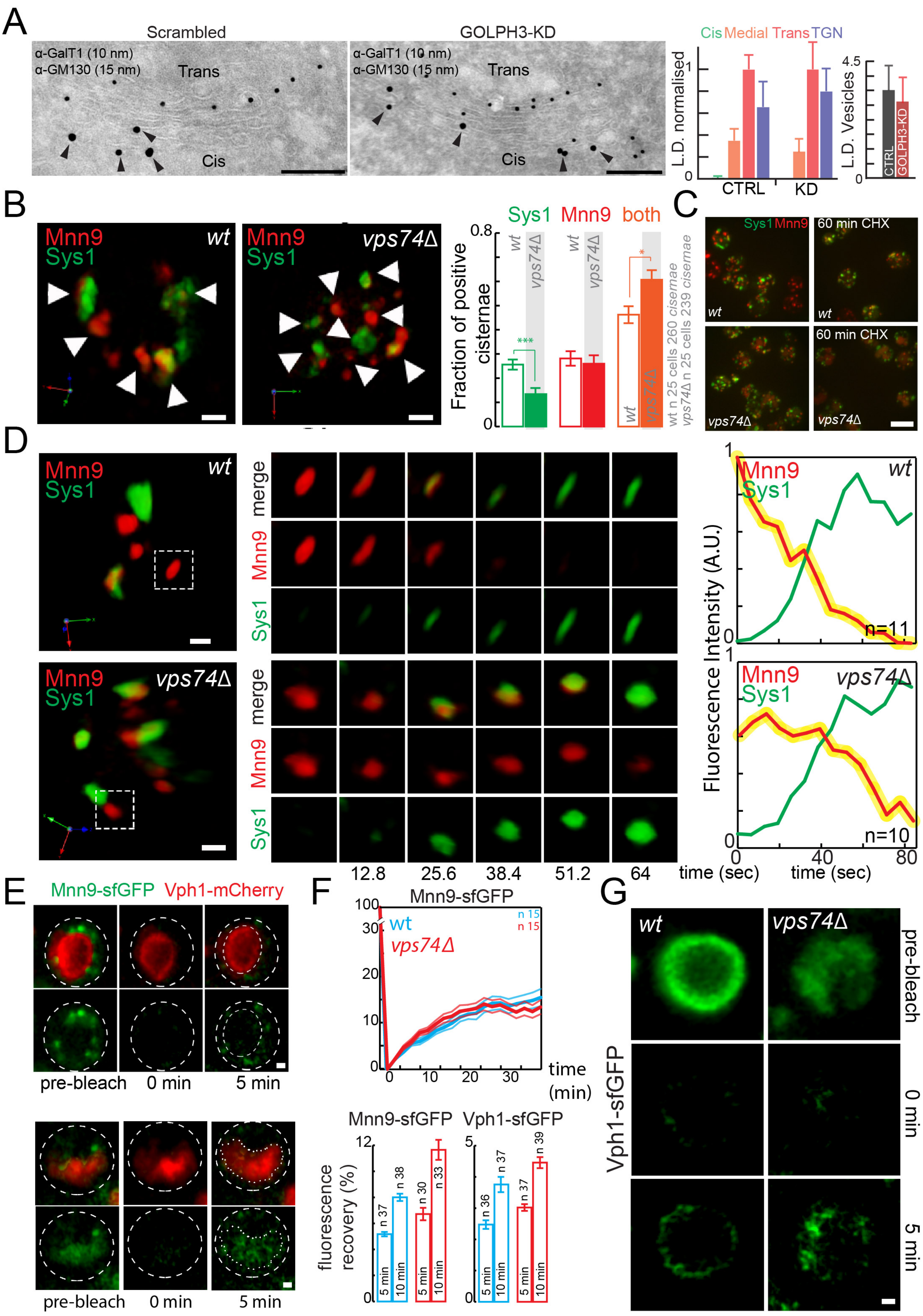

### Supplementary Figure 3

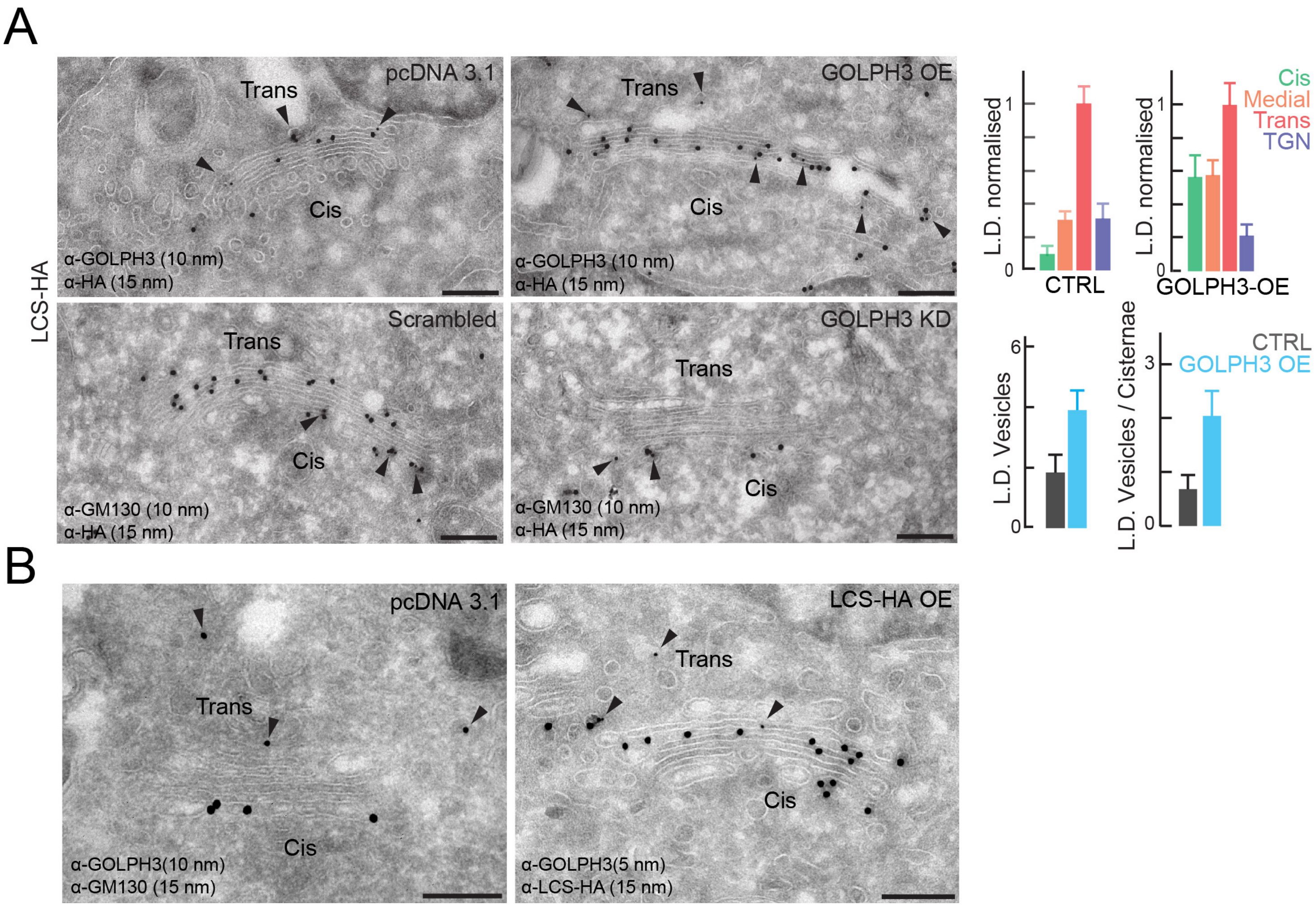

### Supplemetary Figure 4

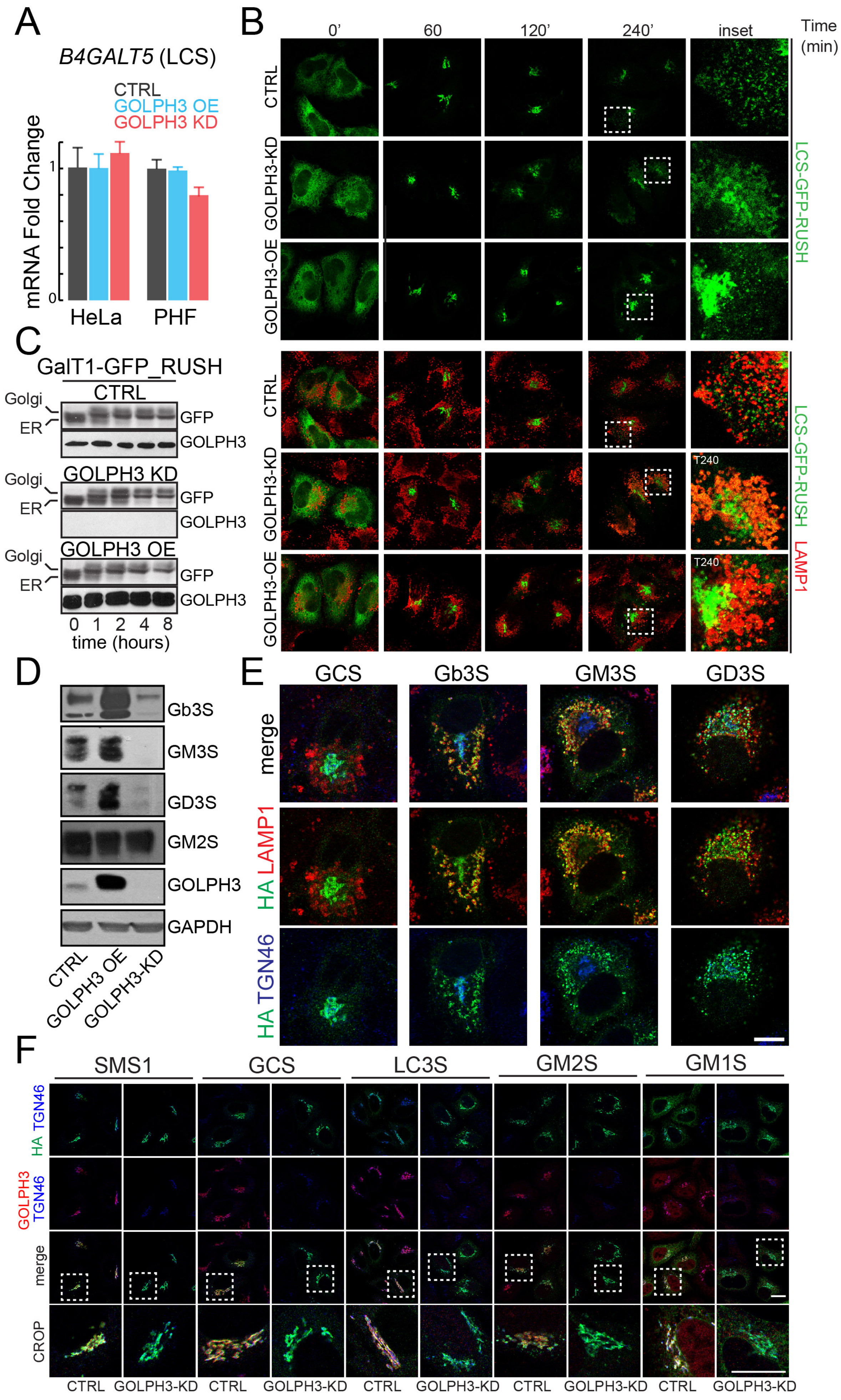

### Supplemetary Figure 5

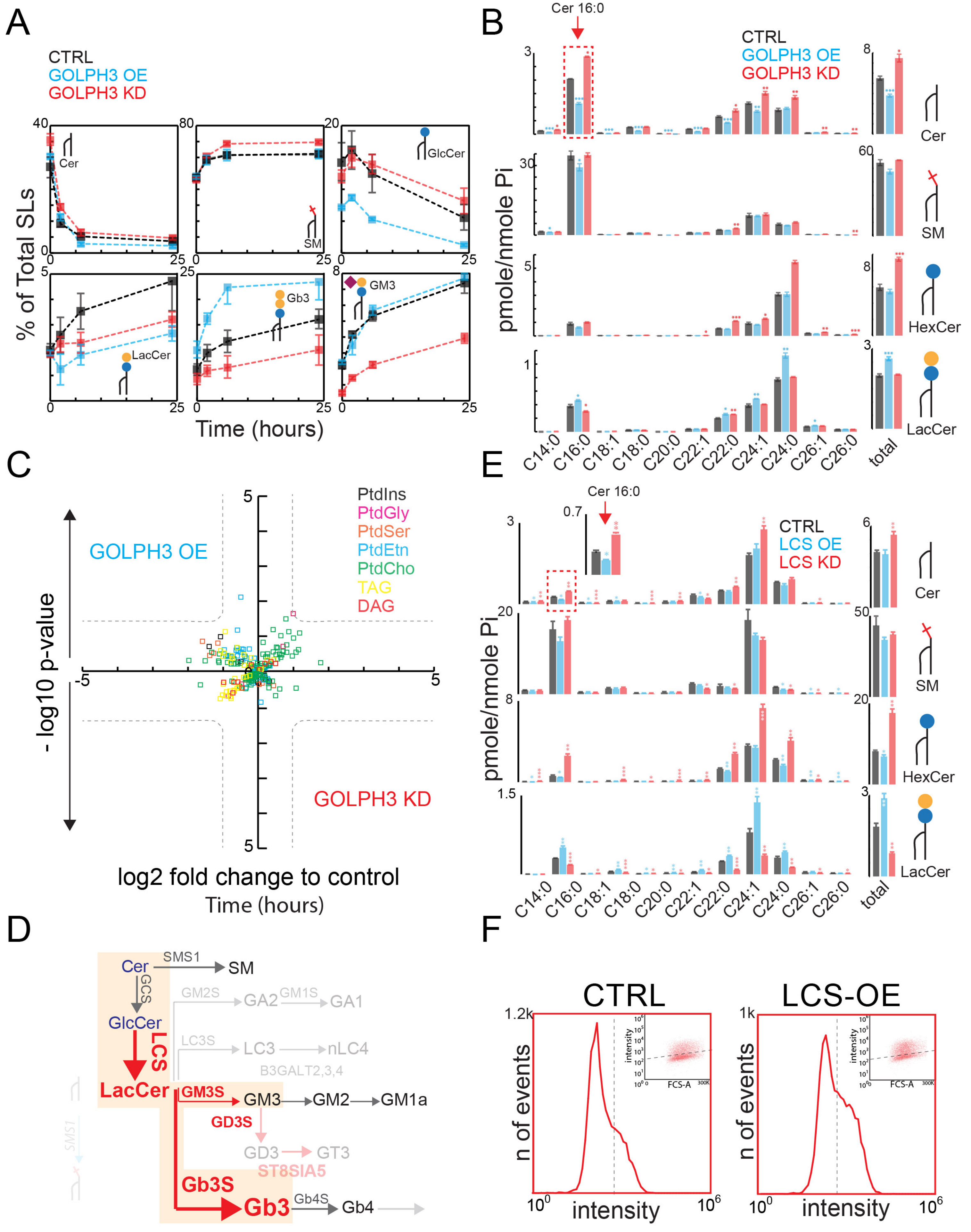

### Supplemetary Figure 6

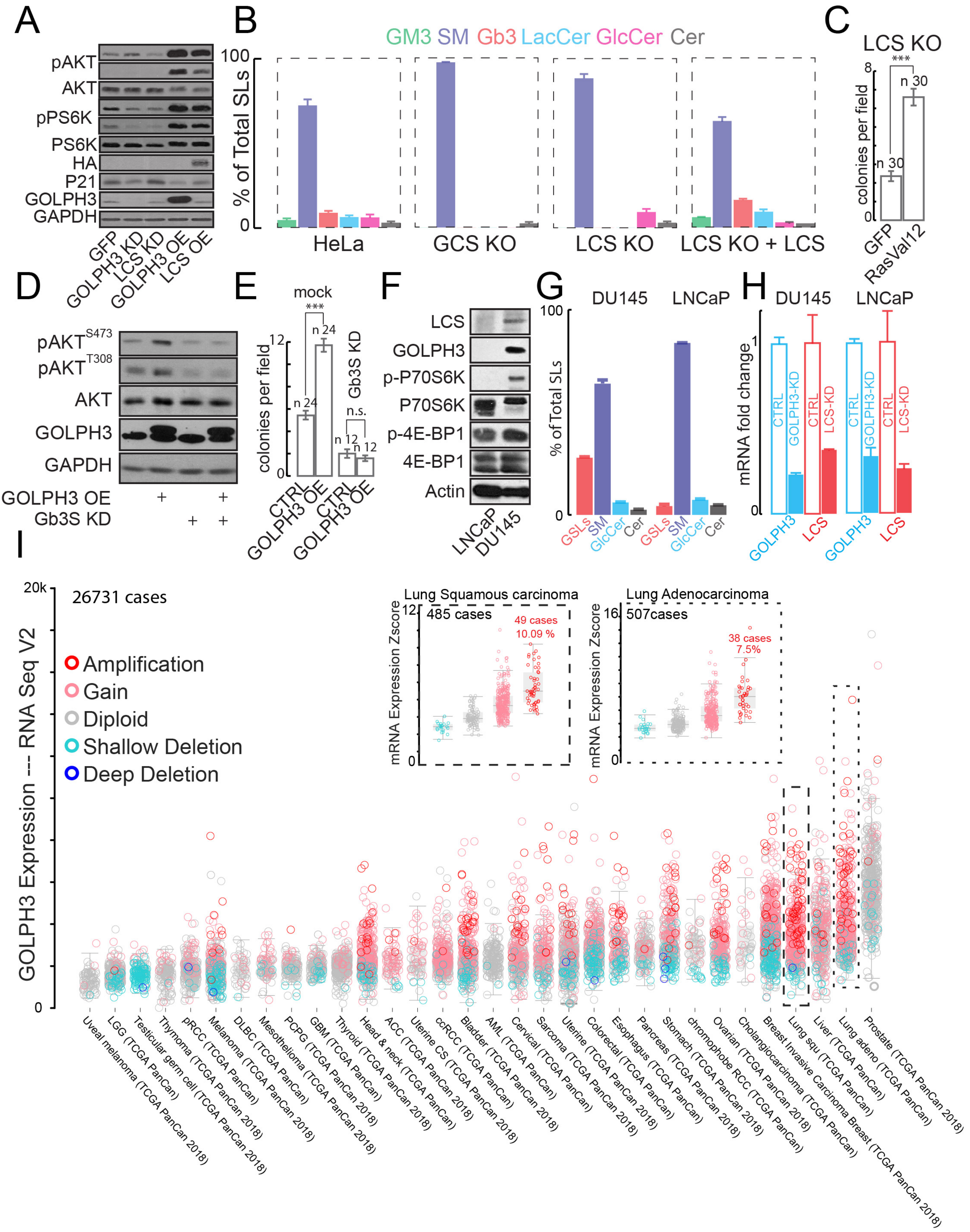
